# Pupil size correlates with near-threshold detection performance irrespective of stimulus colour, eccentricity, or retinal adaptation-state

**DOI:** 10.1101/2025.06.03.657576

**Authors:** Veera Ruuskanen, Sebastiaan Mathôt

**Affiliations:** Department of Experimental Psychology, University of Groningen, The Netherlands

**Keywords:** pupil size, near-threshold detection, retinal adaptation, colour-vision, target-eccentricity

## Abstract

In visual near-threshold detection tasks, larger pre-stimulus pupil size is associated with improved accuracy. However, previous studies have used black and white peripherally presented stimuli, leaving open the question of whether the relationship persists when targets differ in their colour or eccentricity. Here we addressed this question with three experiments that systematically varied the lighting conditions and target properties in a visual near-threshold detection task (data collected in 2022-2024). Light conditions ranged from dark to dim to bright, with the dark condition including a period of dark-adaptation. Possible target colours were blue and red on a black background (dark condition), blue and red on a grey background (dim condition), or yellow and cyan on a white background (bright condition). Possible target eccentricities ranged from parafoveal to peripheral, in a continuous manner (experiment 3) or as two predefined near and far eccentricities (experiments 1 & 2). Across all experiments we show that larger pre-stimulus pupil size is associated with improved performance. This large-pupil advantage is not systematically modulated by the colour or eccentricity of the targets, the illumination of the testing room, or retinal adaptation-state. We conclude that the phenomenon is robust, indicating that pupil size affects vision in a behaviourally relevant manner, regardless of the exact conditions.

**Significance statement:** By showing that pupil size is robustly associated with performance in visual near-threshold detection tasks, we provide evidence for the functional role of endogenous pupil size fluctuations in visual perception.

---

In near-threshold detection tasks, where targets are presented at, or close to, the perceptual threshold, the probability of detection depends on the state of the observer rather than the properties of the stimulus. In the context of visual perception, one aspect of the observer’s state is pupil size, which fluctuates and may influence performance. Previous studies have demonstrated that larger pre-stimulus pupil size is associated with better detection performance (Eberhardt et al., 2022; Mathôt & Ivanov, 2019; Ruuskanen et al., 2025). However, these studies have mainly used black or white peripherally presented stimuli, presented under dim light conditions, leaving open the question of whether the association holds for other types of stimuli and under other light conditions. Yet this is an important question, because the way in which a visual stimulus is processed depends strongly on where in the visual field the stimulus is presented, what colour the stimulus has, and what the adaptation state of the retina is at the moment of stimulus presentation. Furthermore, some evidence suggests that changes in pupil size may lead to changes in the relative activity of the photoreceptors on the retina (Franke et al., 2022). Here we extend previous findings by testing the relationship between pupil size and visual detection performance for different stimulus colours and eccentricities, while also systematically varying the lighting conditions of the experiments.

If visual input is kept constant, the size of the pupil fluctuates as a result of cognitive activity (Ebitz & Moore, 2019; Grujic et al., 2024; Laeng et al., 2012; Mathôt, 2018). These cognitively driven changes in pupil size are believed to result mainly from underlying fluctuations in arousal, due to the link between pupil size and activity in the arousal-regulating locus-coeruleus norepinephrine (LC-NE) system (Bradley et al., 2008; Grujic et al., 2024; Joshi et al., 2016; Joshi & Gold, 2020). Thus, pupil size is commonly used as a marker of arousal and mental effort during cognitive tasks. However, the size of the pupil also affects the amount of light that enters the eye, and how well that light is focused on the retina. Through this route, pupil size affects visual processing and behaviour (Vilotijević & Mathôt, 2023). In other words, pupil size has a bi-directional relationship with higher level cognition and arousal.

One aspect of visual perception in which pupil size plays an important role is optimising vision for detection and discrimination. When the pupil is large, more light is let into the eye, which improves the detection of faint stimuli (Eberhardt et al., 2022; Mathôt & Ivanov, 2019; Ruuskanen et al., 2025; Vilotijević & Mathôt, 2023). This is a robust finding that occurs both when pupil size is manipulated (via changes in background brightness: Eberhardt et al., 2022; Mathôt & Ivanov, 2019) and as a result of spontaneous fluctuations in pupil size (Ruuskanen et al., 2025). Conversely, when the pupil is small, incoming light is focused better, improving discrimination of fine detail (Campbell, 1957; Campbell & Gregory, 1960; Mathôt & Ivanov, 2019; Vilotijević & Mathôt, 2023; Woodhouse, 1975).

In addition to optical effects, detection performance is also affected by arousal. However, while the optical effect manifests itself as a linear relationship between pupil size and performance, the arousal effect is rather inverted-U shaped. This means that performance is best at medium levels of arousal, as opposed to low and high arousal-states (Yerkes & Dodson, 1908). Given the link between pupil size and arousal (Grujic et al., 2024; Joshi & Gold, 2020), this effect is also reflected in the size of the pupil, suggesting that performance is best at medium pupil sizes. Indeed, several studies in both rodents and humans have demonstrated inverted-U shaped relationships between pupil size and performance in tasks related to perceptual decision making with visual, auditory, and tactile stimuli (Beerendonk et al., 2024; de Gee et al., 2024; Doll et al., 2025; McGinley et al., 2015; Podvalny et al., 2021; Schriver et al., 2018).

In addition to factors like pupil size and arousal, perception of visual stimuli is also affected by the properties of the eye, or more specifically, the retina. The retina is composed of two types of photoreceptors (rods and cones) that differ in their overall light sensitivity, their distribution across the retina, and the specific wavelengths of light that they are maximally sensitive to (Barbur & Stockman, 2010; Curcio et al., 1990; Kolb et al., 1995). Firstly, rods are more sensitive than cones, meaning that they require less light to become active. Secondly, rods are more present in the periphery of the retina, while cones are more present in the centre (with the fovea being completely void of rods) (Curcio et al., 1990). Finally, there are three different types of cones that are sensitive to short (peak 420 nm), medium (peak 534 nm), and long (peak 564 nm) wavelengths of light, corresponding to blue, green, and red, respectively. Rods on the other hand are sensitive to wavelengths between short and medium (peak at 498 nm), corresponding to bluish-green light (Barbur & Stockman, 2010; Kolb et al., 1995). Considering these factors, the relationship between pupil size and detection may, in ways that are difficult to predict given our current understanding of the effects of pupil size on vision, vary for stimuli that differentially activate either rods or cones.

Crucial evidence for this possibility comes from a recent study in mice, which showed that changes in pupil size are by themselves sufficient to induce a switch from rod-dominated to cone-dominated vision (Franke et al., 2022). Mice have three types of photoreceptors: the first two types, rods and medium-wavelength (M-)cones, are predominantly sensitive to green light; the third type, short-wavelength (S-)cones, are predominantly sensitive to blue light. In addition, the density of S-cones is highest in the ventral retina, which corresponds to the upper visual field. As a consequence, pupil dilation in mice mainly increased sensitivity to blue light in the upper visual field, and much less so (or not at all) for other colors of light and the lower visual field. There are many differences between the mouse and the human retina, and it is therefore unclear whether and how the results of this study with mice translate to studies with human participants. However, the possibility that the relationship between pupil size and visual sensitivity depends on the location and color of the to-be-detected stimuli is clearly an important one to test.

To investigate this, as well as to further our understanding of how pupil size affects visual processing in general, we conducted three experiments, where participants completed near-threshold detection tasks with varying target colours, eccentricities, and light conditions. We expected to find that, overall, larger pupils are associated with improved detection performance. The shape and strength of this general relationship may vary for different types of targets and different light conditions. However, since this is the first study to systematically address this question, we do not formulate specific hypotheses regarding this.

## Transparency and openness

Details of the methods, data exclusion criteria, and data analysis steps of each experiment can be found below. We did not conduct an a-priori power analysis, instead, sample sizes were determined following the conventions of our lab, whereby we aim for 1600 observations per condition (Brysbaert & Stevens, 2018). This is usually achieved with approximately 30 participants in simple behavioural experiments. For Experiment 3 we aimed for a larger sample size due to the experimental design not allowing for perfect counterbalancing. To further corroborate our chosen sample sizes we have conducted a simulated power analysis using bootstrap samples from another dataset; the details can be found below. Experimental materials, raw data, and analysis scripts are available on the open-science framework (https://osf.io/mzhcx/). None of the experiments were pre-registered.

### Power analysis

The simulated power analysis was performed on bootstrap samples drawn from the dataset described in Ruuskanen and colleagues (2025), where participants completed the same kind of detection task that was used in the current experiments. Due to the nature of our analysis (generalised linear mixed effects modelling) and the fact that this power analysis is based on an existing sample of data, specifying an expected effect size was not possible. Instead, we analysed the power to detect *an effect* (i.e., a slope with an associated p-value < .05) of pupil size on accuracy, using a GLM fitted with participant-specific intercepts and slopes.

To approximate the minimum sample size needed to have sufficient power (80% <) to detect an effect, we tested samples of N=16, N=32, N=64, and N=128, always including 400 trials per subject. For each N, we conducted 1000 simulations by sampling with replacement from our existing dataset, which contained 16 subjects with 840 trials per subject. For each simulation we fitted the GLM and recorded the value of the slope and whether it was significant. Simulations where the model did not converge or resulted in a singular fit were removed. Finally, power at each N was computed as the percentage of iterations where the pupil effect was significant. Estimated power was 70% for N=16, 93% for N=32, 99% for N=64, and nearly 100% for N=128. Based on this analysis we conclude that each experiment described here is sufficiently powered to detect an effect of pupil size on detection accuracy.

## Results overview

An overview of the relationship between pupil size and detection accuracy (defined as the combined proportion of hits and correct rejections) in each experiment is depicted in Figure 1. This relationship was significant (*p* < 0.001; analysed using generalised linear mixed models) in all datasets except for the dark condition of Experiment 3 (*p* = 0.06), although also in this condition the effect was numerically in the same direction Further details of the methods and results of each experiment are described below, but the main finding across all experiments is: larger pupils are robustly associated with increased detection performance across a wide range of different stimulus types and conditions.

**Figure 1:**
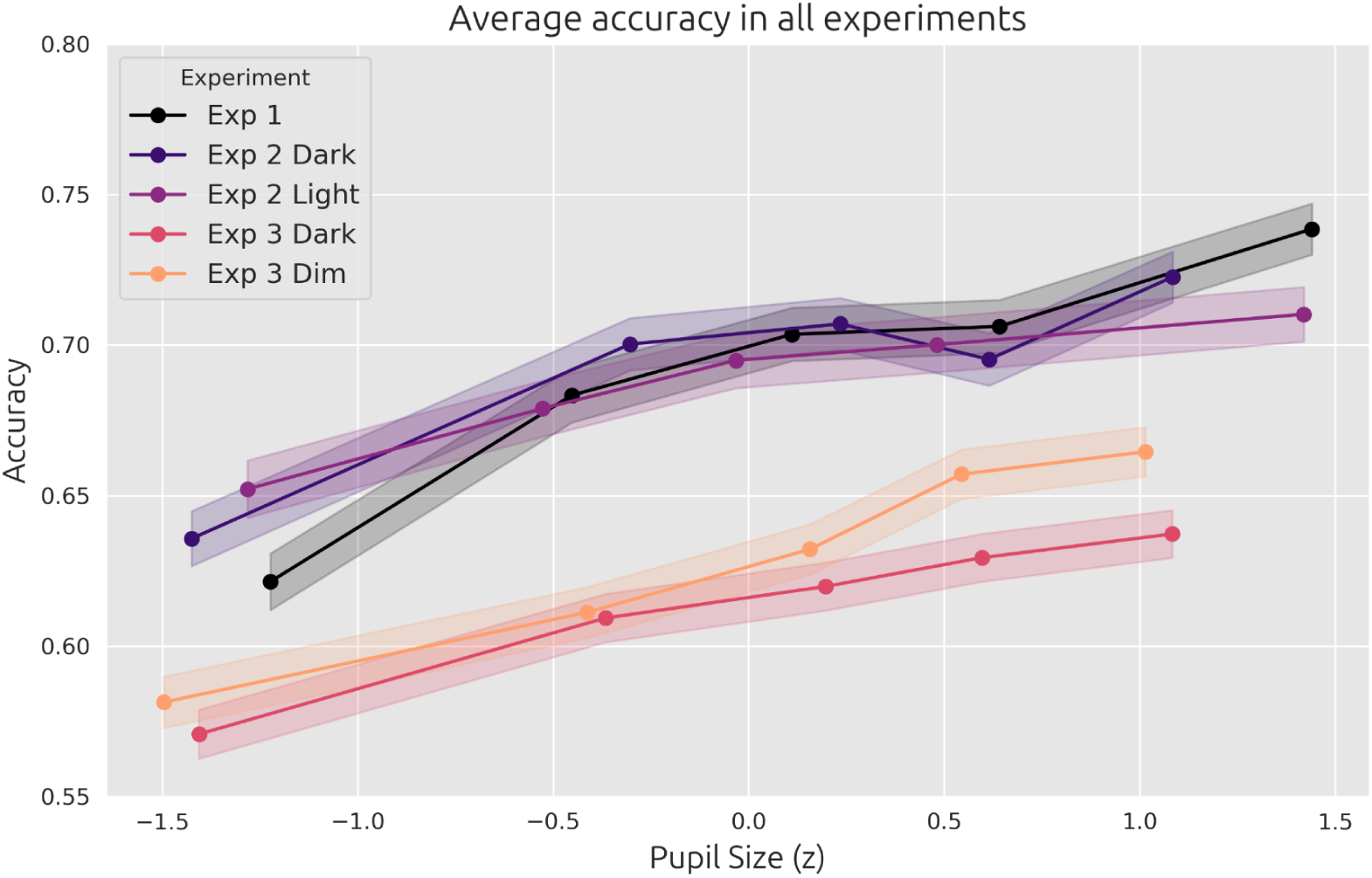
Accuracy as a function of pre-stimulus pupil size in all experiments. Lines represent separate datasets from experiments 1, 2, and 3. For each experiment, pupil size is normalised within subjects and binned into 5 bins for plotting. Dots on the lines represent the average of each bin. Accuracy refers to the combined proportion of hits and correct rejections. The shaded area around the lines represents the standard error.

## Experiment 1

### Introduction

In Experiment 1, we set out to investigate how stimuli of different colours and eccentricities interact with pupil size in determining detection performance under constant light conditions. The two colours (blue and red) and two eccentricities (parafoveal and peripheral) were chosen in order to maximally differentiate between stimuli that activate either the rods or the cones. Specifically, due to the wavelength of light that rods are maximally sensitive to, they are most sensitive to blue light, but less sensitive to red light. Thus, the detection of red stimuli is assumed to rely more on the activation of cones rather than rods. (In this experiment, stimuli were presented on a gray background, which complicates the relationship between stimulus color and photoreceptor activation. We will return to this point in later experiments.) Furthermore, due to the higher density of rods in the periphery of the retina, the detection of especially peripherally presented blue targets is assumed to rely more on the activation of rods rather than cones.

## Method

### Participants

30 participants with normal (colour) vision participated in the experiment. All participants were undergraduate students at the University of Groningen and received partial course credit in exchange for participation. The data was collected in the year 2022. No participants were excluded. All participants received information about the experiment and provided written informed consent prior to participation. Based on criteria developed by the EC-BSS at the University of Groningen, the study was exempt from full ethical review (study code: PSY-2223-S-0051).

### Experimental design

#### Software, apparatus, and data acquisition

The experimental task was programmed in OpenSesame (version 4.0.10, *Melodramatic Milgram*) (Mathôt et al., 2012) and presented on an 27-inch PROLITE G2773HBS-GB1 (EOL) monitor running at a refresh rate of 60 Hz and 1920 × 1080 pixels resolution. Pupil data was recorded with an Eyelink 1000 eye tracker (SR Research Ltd., 2022) at a sampling frequency of 1000 Hz. An initial 9-point calibration procedure was performed at the beginning of the experiment and a drift correction was performed at the beginning of each trial. A chinrest was used to keep viewing distance fixed at approximately 60 cm. The experiment was conducted in a dimly lit room (illumination as measured at the chinrest 6 LUX).

#### Detection task and procedure

Participants completed a near-threshold visual detection task (illustrated in Figure 2). The task was presented on a grey (RGB = 128, 128, 128) background with a luminance of 30.13 cd/m^2^ (CIE coordinates: x ≈ 0.314, y ≈ 0.324). A circular grey fixation dot (RGB = 77, 77, 77) with a size of 0.44° of visual angle (15 px) was maintained in the centre of the screen throughout the experiment.

**Figure 2:**
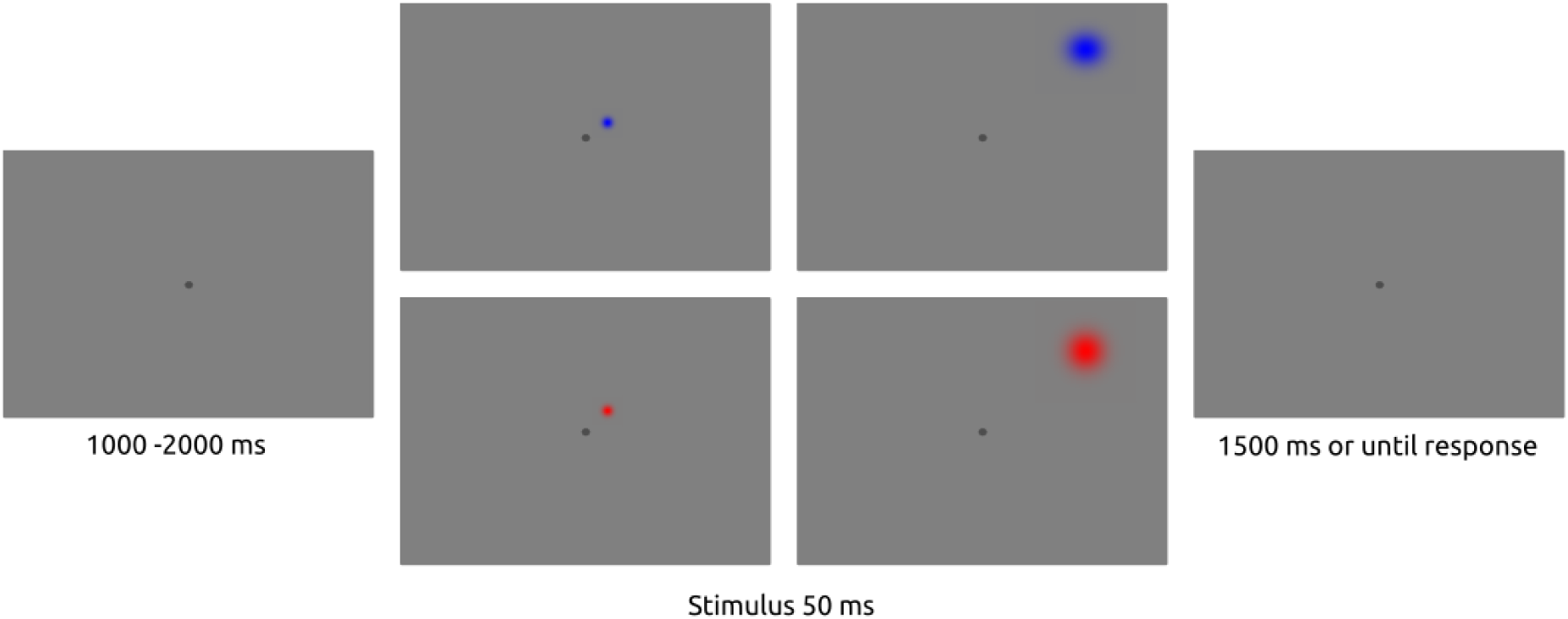
The detection task in Experiment 1. A stimulus was flashed on 50% of the trials. This illustration is not to scale and the targets are brighter than in the actual experiment.

The detection targets consisted of red (RGB = 255, 0, 0) and blue (RBG = 0, 255, 0) luminance patches enveloped in a Gaussian envelope. The luminance of the stimuli at full contrast was 25.39 cd/m^2^ for red (CIE coordinates: x ≈ 0.654, y ≈ 0.342) and 7.43 cd/m^2^ for blue (CIE coordinates: x ≈ 0.151, y ≈ 0.057). However, the starting contrast was 40% for blue targets and 30% for red targets (determined at the authors’ discretion) and was further adjusted with a staircase procedure to keep overall accuracy fixed at 65% thus leading to a lower stimulus-luminance during the task. The staircase worked with a 1-up 3-down procedure, whereby the stimulus opacity was increased (by 10% of the current value) after each incorrect response and decreased (by 10% of the current value) after three consecutive correct responses.

Stimuli were presented on an invisible circle at either a near or a far eccentricity. ‘Near’ targets were presented parafoveally at 0.58° of visual angle (20 px), while ‘far’ targets were presented in the periphery at 8.71° of visual angle (300 px). Exact stimulus location was determined by drawing a random angle between 0° and 360° and transforming it to x and y coordinates considering the given eccentricity. The size of the stimuli was adjusted according to the cortical magnification formula (1) such that the ‘near’ targets were smaller than the ‘far’ targets (taken from Carrasco & Frieder, 1997).

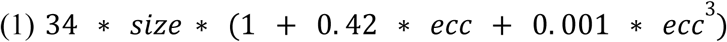

Each trial started with a variable start time drawn from a uniform distribution between 1 and 2 seconds. Next, on half of the trials a target was flashed for 50 ms. The target-present trials contained an even distribution of possible combinations of colour and eccentricity, creating 4 conditions: blue-near, blue-far, red-near, and red-far. Following target onset, each trial lasted for another 1.5 seconds, making total trial length between 2.5 and 3.5 seconds. Since the fixation dot was maintained on the screen throughout the experiment, the participants were not aware of the beginning and end of each trial, thus perceiving the task as a continuous stream. Participants were asked to report whenever they detected a target by pressing the spacebar. Upon keypress the fixation dot changed into a square for 10ms, creating a visual transient intended to indicate that a response was recorded.

The task consisted of a practice phase and experimental phase. The practice phase lasted for 16 trials, during which the targets were slightly brighter (15% contrast) than during the experimental phase, to familiarise the participants with the task. Participants received task instructions both verbally and on the screen prior to practice. The experimental phase consisted of 10 blocks of 64 trials, separated by self-paced breaks. The staircase procedure was implemented during the first 3 blocks and these were excluded from the analysis. Thus, 448 experimental trials were included in the analysis.

The whole experimental session, including informed consent, eye tracker calibration, task performance, and debriefing, took approximately 1 hour and 15 minutes per participant.

#### Data processing and analysis

The pupil data was processed using the Eyelinkparser (Mathôt & Vilotijević, 2022) package in Python. The data was downsampled to 100 Hz and blinks were interpolated. Pupil size was then transformed from arbitrary units to millimetres. Trials with biologically implausible pupil sizes (smaller than 2 mm or larger than 8 mm) were removed. For the analysis, the data was cropped to 1 second before stimulus presentation (or to 1 second prior to when a stimulus *would have been presented* on stimulus absent trials), the average pupil size in this period was extracted, and then z-scored for each participant individually, across blocks.

To examine interactions between pupil size and target colour and eccentricity, two generalised mixed linear models (GLM) were constructed using single-trial data. The first model included z-scored pupil size, target colour, target eccentricity, and all possible interactions as fixed effects, participant-specific slopes and intercepts as random effects and accuracy as the dependent variable. To account for a possible non-linear relationship between pupil size and detection, the second model also included squared pupil size, but was otherwise identical to the first. The models were compared with the Akaike information criterion (AIC), which was lower for the first (simpler) model. Thus, the results of Model 1 are reported here. We do not report effect size estimates because deriving meaningful estimates for GLMs is non-trivial. We do report coefficients and their significance.

While we will report main effects of various stimulus features, in this case colour and eccentricity, these main effects are not of theoretical interest, because they simply reflect that difficulty is not perfectly controlled across these feature dimensions. What is of theoretical interest, however, is whether these stimulus stimulus characteristics interact with pupil size.

## Results

### Descriptive statistics

The distribution of recorded (z-scored) pupil sizes and average pupil size over time are depicted in Figure 3. On average pupil size was larger after breaks (as indicated by dashed lines in Fig 3b) and then decreased over the course of the block. Despite this systematic variation we did not remove any trials from the analysis to avoid excluding genuine (arousal-driven) fluctuations in pupil size. However, in the Supplementary Materials we report the details of a control analysis where time on task during each block is regressed out of the pupil signal. This does not change the pattern of results, and thus, here we report the results of the analysis using the true recorded pupil size. Average accuracy across the task and in each condition (i.e., combination of target colour and eccentricity) can be found in Table 1.

**Figure 3:**
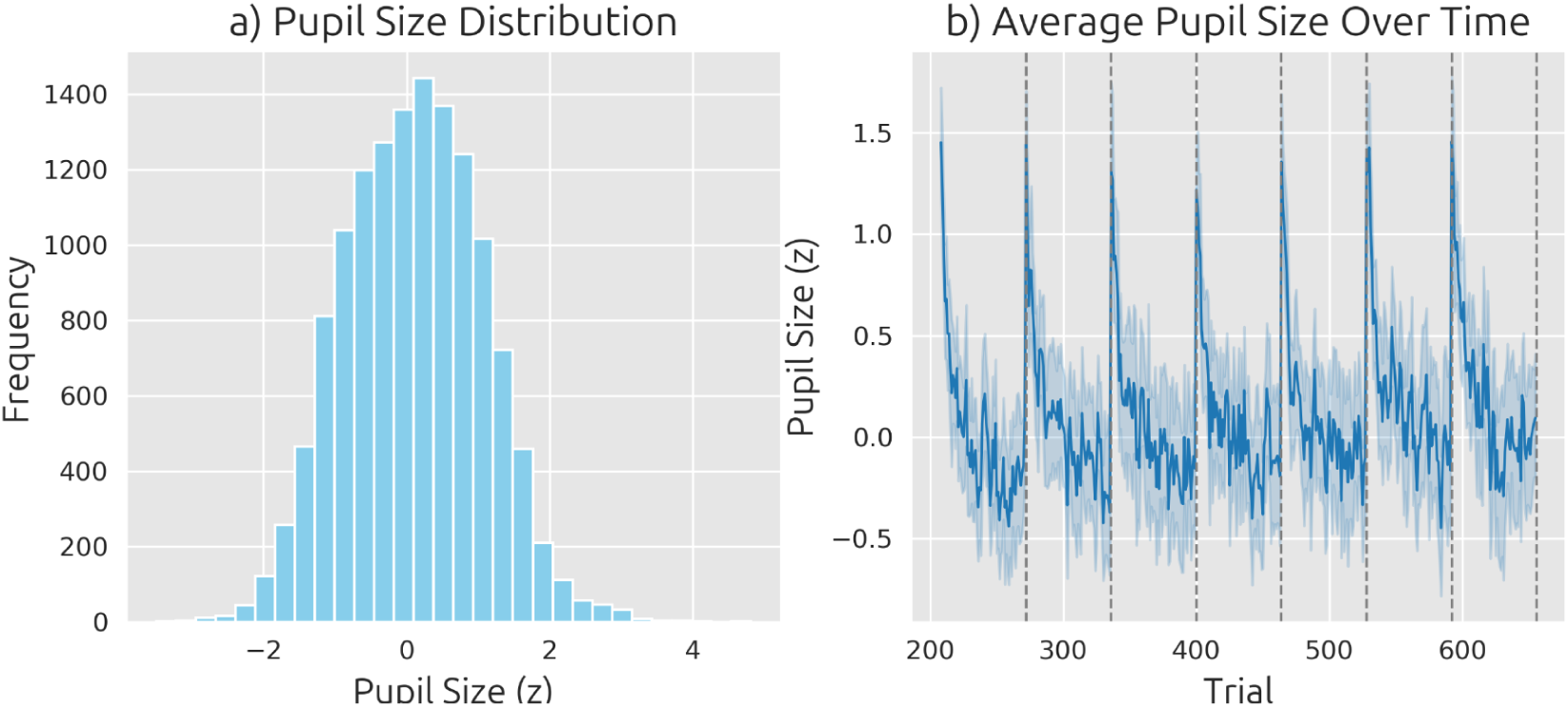
Pupil sizes recorded in Experiment 1 (normalised). a) Distribution of all recorded pupil sizes. b) Pupil size over time. Dashed vertical lines denote breaks. The shaded area around the line represents a bootstrap 95% confidence interval.

**Table 1.** Accuracy in each condition of Experiment 1.

| Condition | M (%) | SD |
| --- | --- | --- |
| All | 68.99 | 0.46 |
| Blue-Far | 69.76 | 0.46 |
| Blue-Near | 67.95 | 0.47 |
| Red-Far | 73.42 | 0.44 |
| Red-Near | 64.82 | 0.48 |

### Main analysis

The GLM showed significant main effects of pupil size (*b* = 0.24, *p* < .001), whereby larger pupils improved detection, and of eccentricity (*b* = 0.13, *p* = .008), whereby peripheral targets were detected better. Further, there was a significant interaction between colour and eccentricity (*b* = 0.08, *p* < .001), whereby red and peripheral targets were detected best. Crucially, however, pupil size did not significantly interact with either colour (*b* =-0.01, *p* = .751) or eccentricity (*b* = -0.00, *p* = .940). The results are depicted in Figure 4.

**Figure 4:**
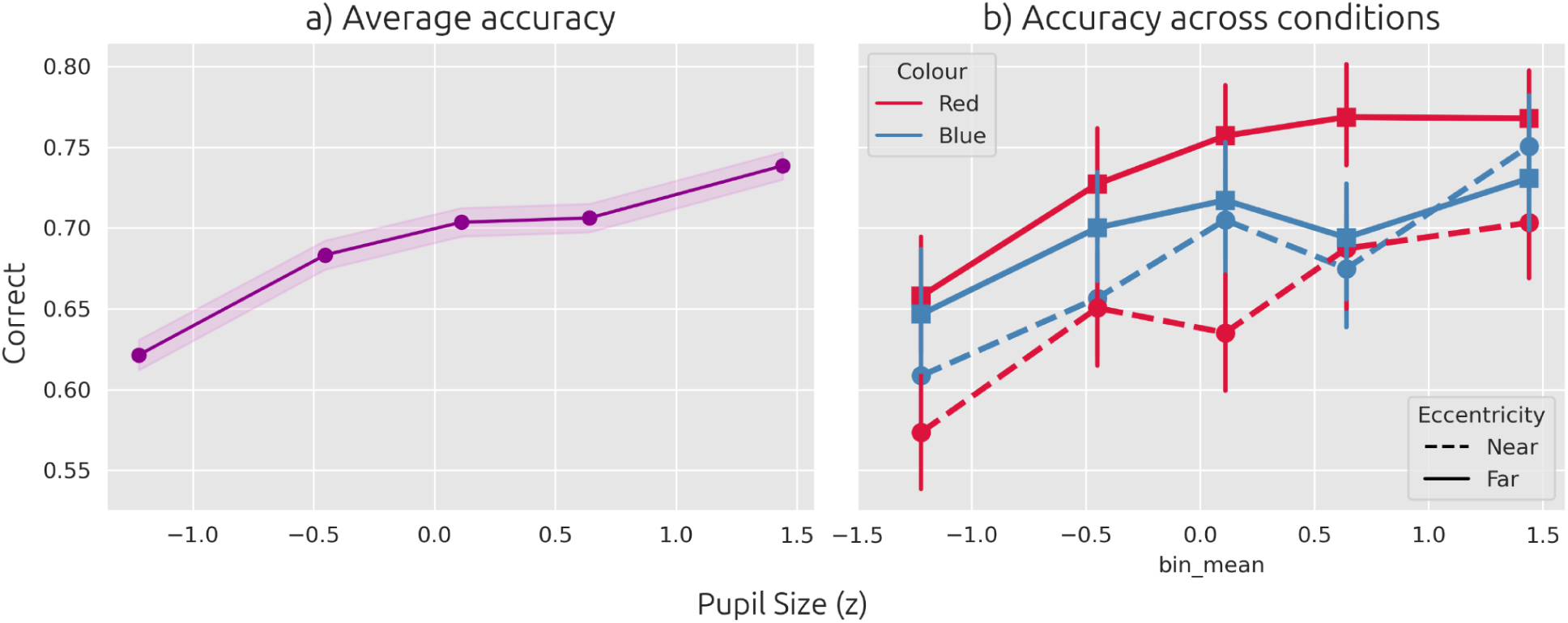
Accuracy as a function of pre-stimulus pupil size in Experiment 1. Pupil size is normalised within participants. Accuracy here refers to hits and correct rejections. A) Overall accuracy. The shaded area around the line represents the standard error. B) Accuracy divided into conditions. The error bars represent a bootstrap 95% confidence interval around the mean.

## Discussion

In experiment 1 we investigated the association between pupil size and detection of visual targets with varying colours (blue and red against a gray background) and eccentricities (parafoveal and peripheral). We found that performance improves with increasing pupil size, replicating previous results (Eberhardt et al., 2022; Mathôt & Ivanov, 2019; Ruuskanen et al., 2025). Further (but as mentioned not of theoretical interest), peripheral targets were overall detected better than parafoveal targets, and performance was best for red and peripheral targets. Importantly, however, we did not observe interactions between pupil size and either colour or eccentricity, meaning that large pupils benefited detection performance regardless of these target-properties. These results do not support the proposition that pupil dilation results in a switch from rod-dominated to cone-dominated vision in humans.

There are some limitations in the study design which makes it difficult to draw firm conclusions. Firstly, adding a target of a certain colour onto a grey background does not only change the absolute amount of that colour being projected on the retina, but also affects the relative amounts of other colours present. Specifically, when a red stimulus is presented on a gray background, not only does the amount of red increase, the amount of blue and green decreases as well. This means that detection performance of red stimuli against a grey background is not a pure measure of sensitivity to red. (And the same argument applies to blue.)

## Experiment 2

### Introduction

In Experiment 2, we adjusted the design to replicate and extend Experiment 1. The aim was again to investigate how stimuli of different colours and eccentricities interact with pupil size in determining detection performance. Furthermore, with the aim of investigating how the adaptation state of the retina interacts with pupil size in determining detection performance and to promote reliance on one photoreceptor class over the other, the task was conducted under two opposing light conditions (dark and bright; promoting scotopic and phototopic vision, respectively). For the dark condition, the colours (blue and red) and eccentricities (parafoveal and peripheral) of the stimuli were kept the same as in Experiment 1, but the background colour was changed to black, so that the presentation of a red stimulus only changed the amount of red. In the bright condition the colours were reversed, with a white background and cyan and yellow targets. Cyan and yellow were chosen due to the white background. White contains an equal amount of all the primary colours, and thus, to activate one colour channel over others, colour must be taken away rather than added. Removing all red makes cyan and removing all blue makes yellow.

### Method

Experiment 2 consisted of two experimental sessions: a dark session and a bright session and the same sample of participants participated in both. The order of experimental sessions was counterbalanced across participants. For both sessions, the software, apparatus, data acquisition procedure and task design were largely the same as in experiment 1. Thus, in this section we will only describe in detail the aspects that differed: the sample of participants, the light conditions, and the colours of the targets and background in the task.

#### Participants

A total of 31 participants with normal (colour) vision participated in the experiment but two participants only completed the dark session. All participants were undergraduate students at the University of Groningen and received partial course credit in exchange for participation. The data was collected in the year 2023. No participants were excluded. All participants received information about the experiment and provided written informed consent prior to participation. Based on criteria developed by the EC-BSS at the University of Groningen, the study was exempt from full ethical review (study code: PSY-2223-S-0434).

#### Experimental design

##### Dark session

In the dark session, the lights in the testing room were turned off (illuminance at chinrest 0 LUX) and potentially distracting light sources - like the power button of the monitor - were covered. Before starting the task, participants underwent 10 minutes of dark-adaptation.

The detection task was presented on a black background (RGB = 0, 0, 0) with a luminance of 0.14 cd/m^2^ (CIE coordinates: x ≈ 0.216, y ≈ 0.299). The to-be-detected targets were the same red and blue luminance patches described above. The size of the targets was adjusted with a cortical magnification formula, as described in Experiment 1. The task is illustrated in Figure 5.

**Figure 5:**
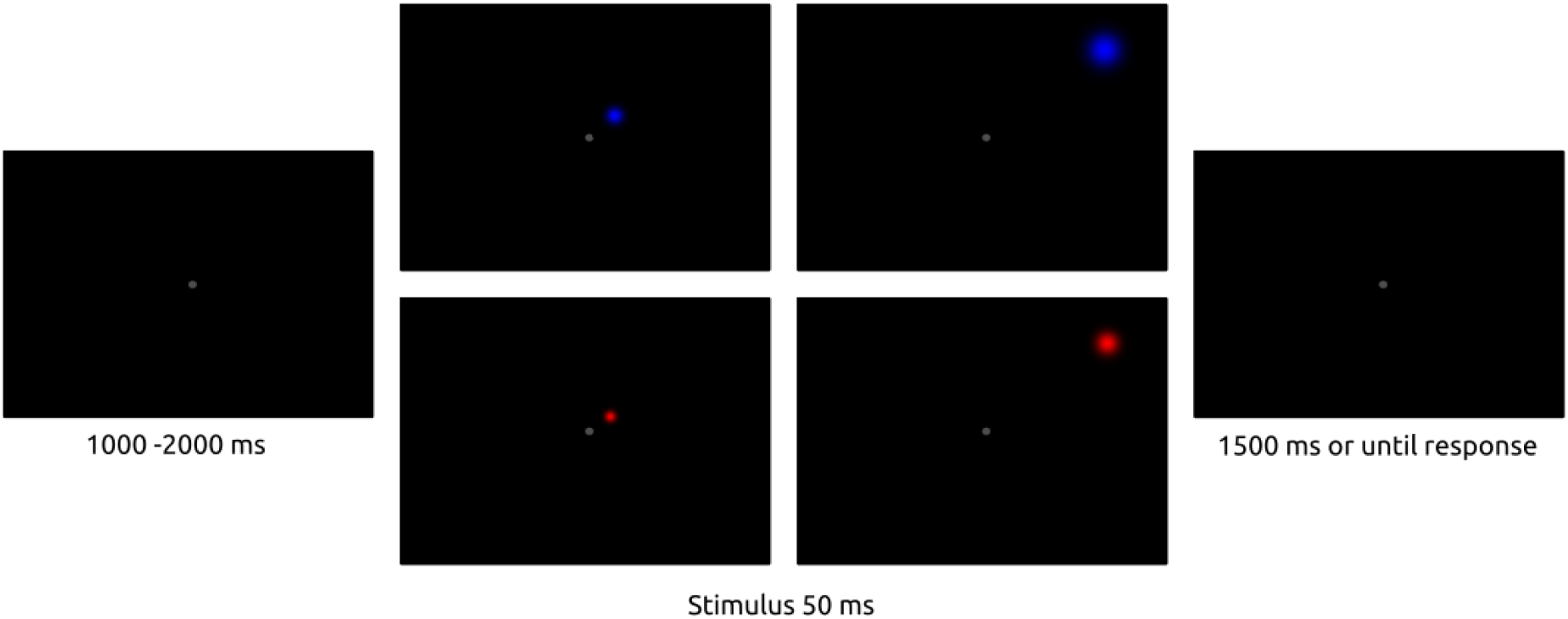
The detection task in the dark session of Experiment 2. A stimulus was flashed on 50% of the trials. This illustration is not in scale and the targets are brighter than in the actual experiment.

The same data processing and analysis pipeline was applied as in Experiment 1 (see above). The simpler model (excluding a pupil squared term) had a lower AIC. Therefore, the results of this model will be reported here.

##### Bright session

In the bright session, the lights in the testing room were turned on as bright as possible (illuminance at chinrest 85 LUX).

The detection task was presented on a white background (RGB = 255, 255, 255) with a luminance of 111.64 cd/m^2^ (CIE coordinates: x ≈ 0.322, y ≈ 0.337). The to-be-detected targets were yellow (RGB = 255, 255, 0) and cyan (RGB = 0, 255, 255) luminance patches, enveloped in a Gaussian envelope. The luminance of the targets at full contrast was 95.3 cd/m^2^ for yellow (CIE coordinates: x ≈ 0.441, y ≈ 0.533) and 77.97 cd/m2 for cyan (CIE coordinates: x ≈ 0.225, y ≈ 0.336). The size of the targets was adjusted with a cortical magnification formula, as described in Experiment 1. The task is illustrated in Figure 6.

**Figure 6:**
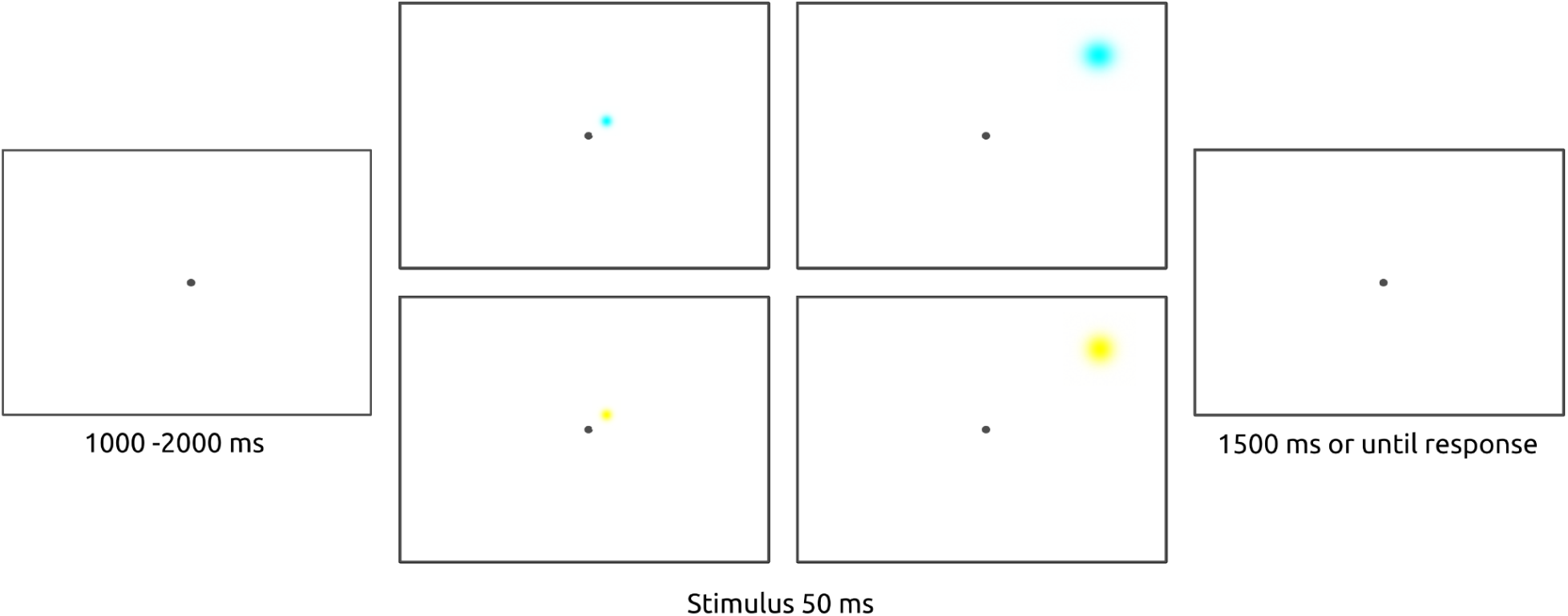
The detection task in the light session of Experiment 2. A stimulus was flashed on 50% of the trials. This illustration is not in scale and the targets are brighter than in the actual experiment.

The same data processing and analysis pipeline was applied as in Experiment 1 (see above). The simpler model (not including a pupil squared term) had a lower AIC. Therefore, the results of this model will be reported here.

## Results

### Dark

#### Descriptive statistics

The distribution of recorded (z-scored) pupil sizes and average pupil size over time are depicted in Figure 7. On average pupil size was larger after breaks (as indicated by dashed lines in Fig 7b) and then decreased over the course of the block. Despite this systematic variation we did not remove any trials from the analysis to avoid excluding genuine (arousal-driven) fluctuations in pupil size. However, in the Supplementary Materials we report the details of a control analysis where time on task during each block is regressed out of the pupil signal. This does not change the pattern of results, and thus, here we report the results of the analysis using the true recorded pupil size. Average accuracy across the task and in each condition (i.e., combination of target colour and eccentricity) can be found in Table 2.

**Figure 7:**
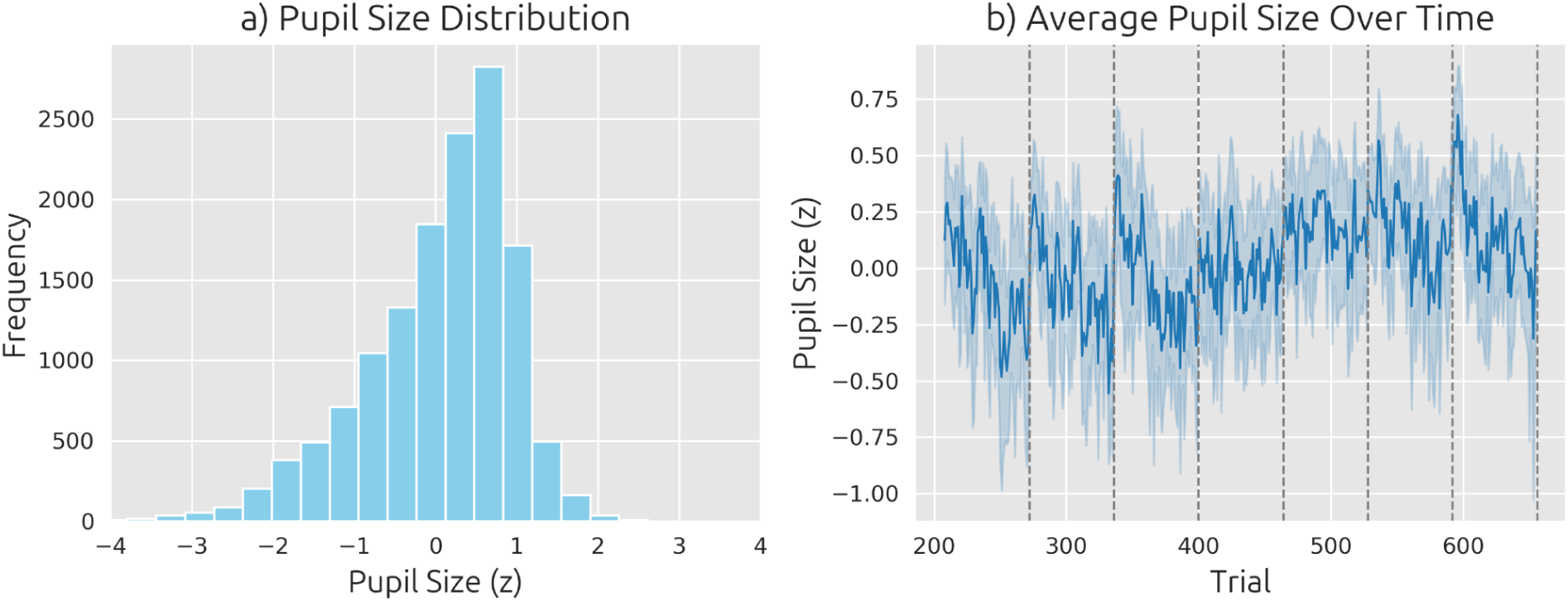
Pupil sizes recorded in the dark condition of Experiment 2 (normalised). a) Distribution of all recorded pupil sizes. b) Pupil size over time. Dashed vertical lines denote breaks. The shaded area around the line represents a bootstrap 95% confidence interval.

**Table 2.**
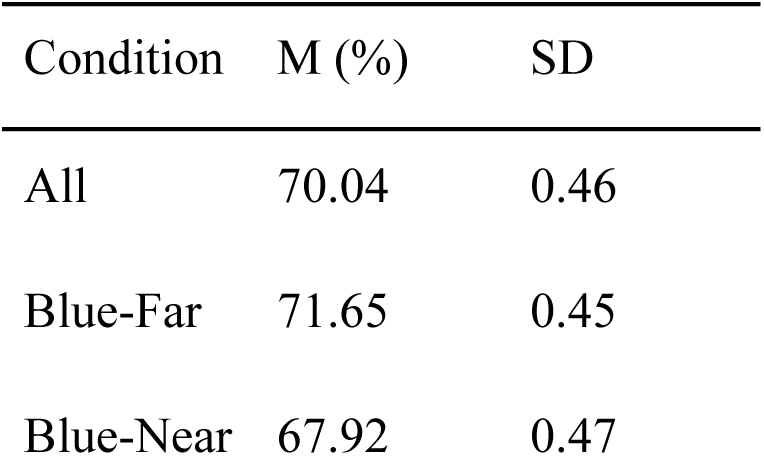
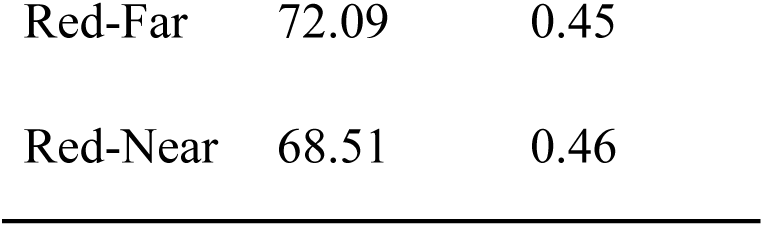
Accuracy in each condition of the dark session of Experiment 2.

| Condition | M (%) | SD |
| --- | --- | --- |
| All | 70.04 | 0.46 |
| Blue-Far | 71.65 | 0.45 |
| Blue-Near | 67.92 | 0.47 |
| Red-Far | 72.09 | 0.45 |
| Red-Near | 68.51 | 0.46 |

#### Main analysis

The GLM showed significant main effects of pupil size (b = 0.14, p < .001), whereby larger pupils improved detection. Pupil size did not significantly interact with either colour (*b* = 0.02, *p* = .301) or eccentricity (*b* = 0.00, *p* = .95). The results are depicted in Figure 8.

**Figure 8:**
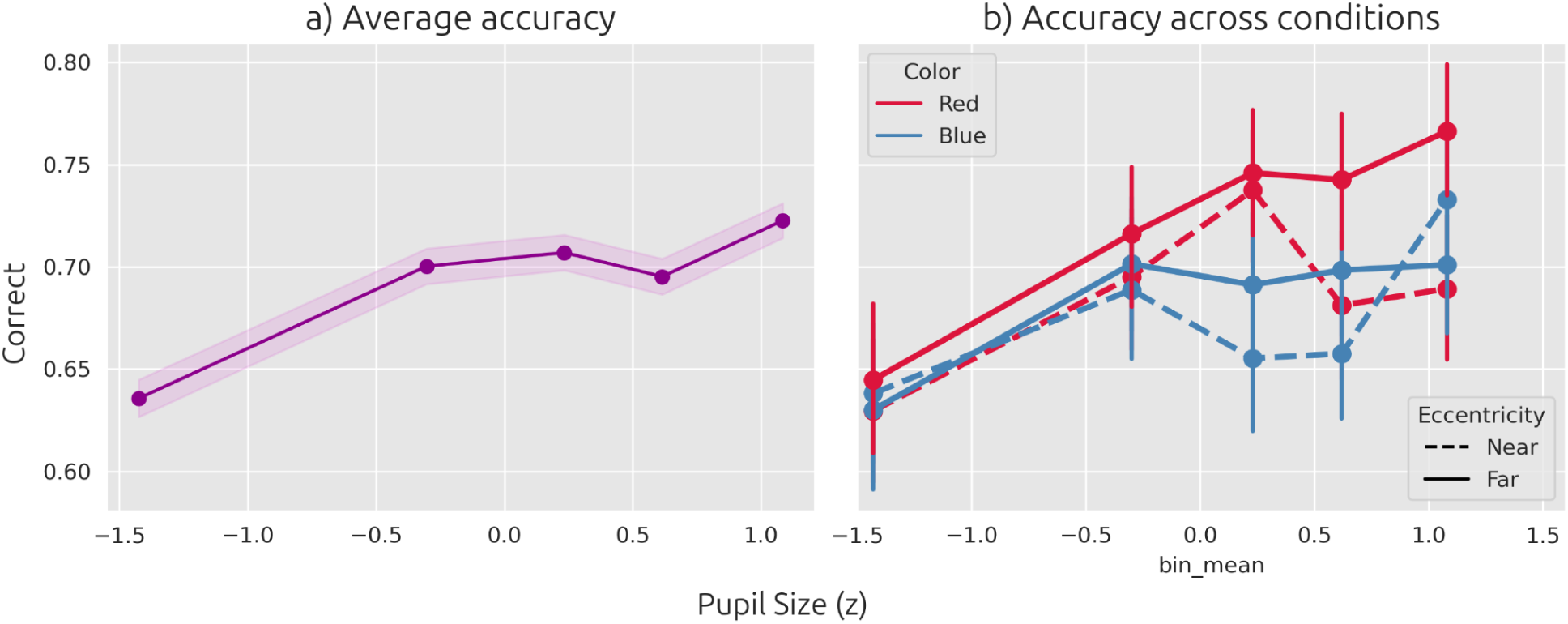
Accuracy as a function of pre-stimulus pupil size in the dark condition of Experiment 2. Pupil size is normalised within participants. Accuracy here refers to hits and correct rejections. A) Overall accuracy. The shaded area around the line represents the standard error. B) Accuracy divided into conditions. The error bars represent a bootstrap 95% confidence interval around the mean.

### Bright

#### Descriptive statistics

The distribution of recorded (z-scored) pupil sizes and average pupil size over time are depicted in Figure 9. On average pupil size was larger after breaks (as indicated by dashed lines in Fig 9b) and then decreased over the course of the block. Despite this systematic variation we did not remove any trials from the analysis to avoid excluding genuine (arousal-driven) fluctuations in pupil size. However, in the Supplementary Materials we report the details of a control analysis where time on task during each block is regressed out of the pupil signal. This does not change the pattern of results, and thus, here we report the results of the analysis using the true recorded pupil size. Average accuracy across the task and in each condition (i.e., combination of target colour and eccentricity) can be found in Table 3.

**Figure 9:**
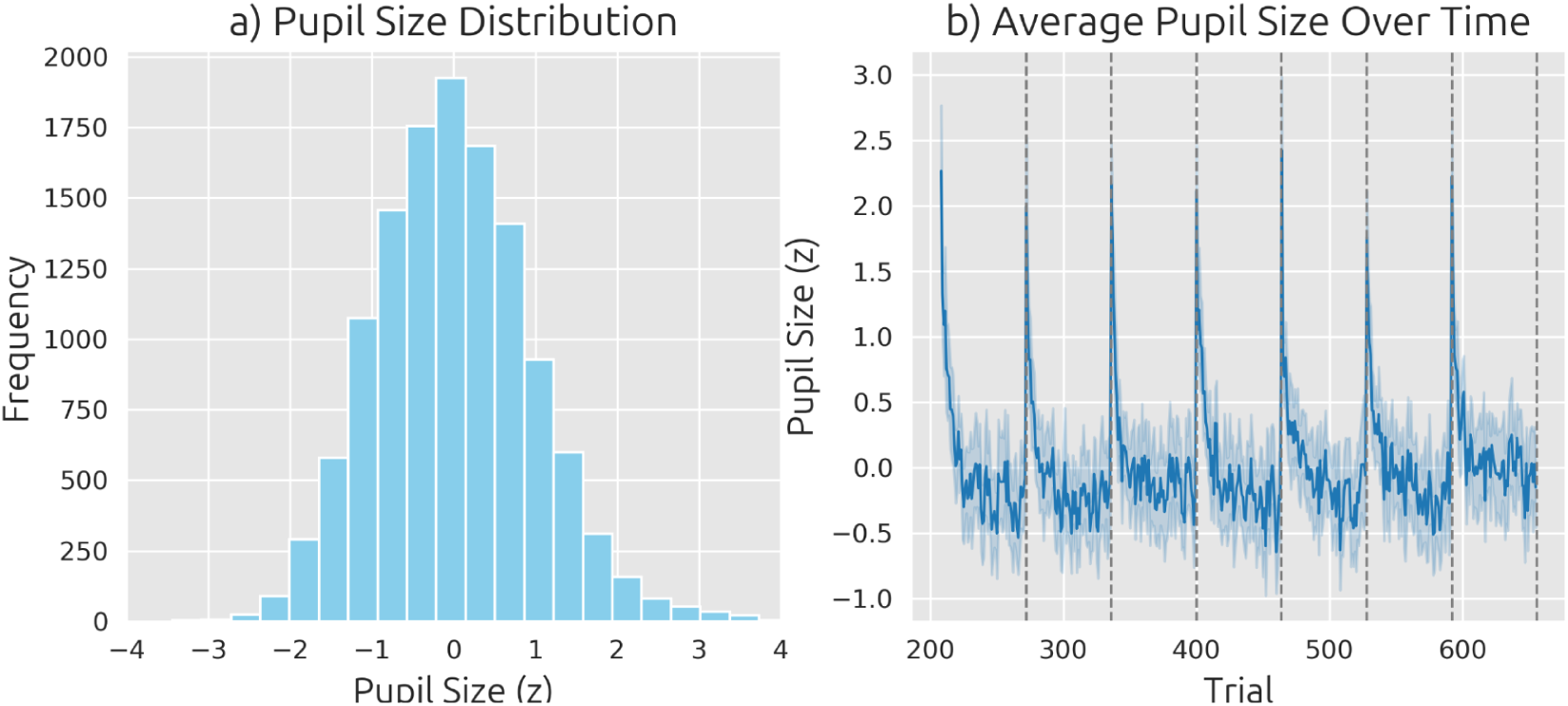
Pupil sizes recorded in the bright condition of Experiment 2 (normalised). a) Distribution of all recorded pupil sizes. b) Pupil size over time. Dashed vertical lines denote breaks. The shaded area around the line represents a bootstrap 95% confidence interval.

**Table 3.**
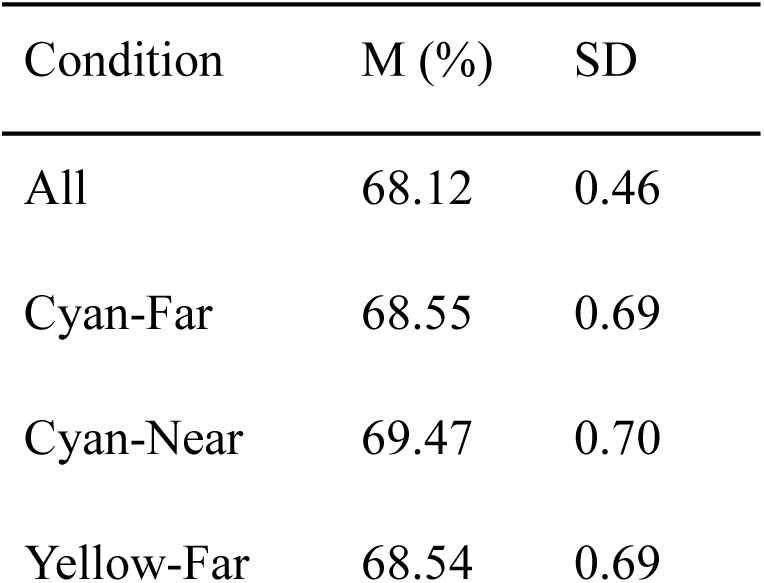
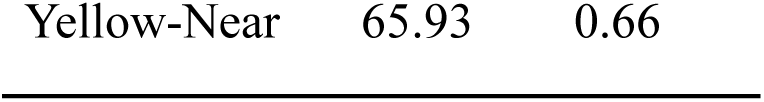
Accuracy in each condition of the bright session of Experiment 2.

#### Main analysis

The GLM showed a significant main effect of pupil size (b = 0.08, p < .001), whereby larger pupils improved detection. However, pupil size did not significantly interact with either colour (*b* = 0.01, *p* = .571) or eccentricity (*b* = 0.01, *p* = .554). The results are depicted in figure 10.

**Figure 10:**
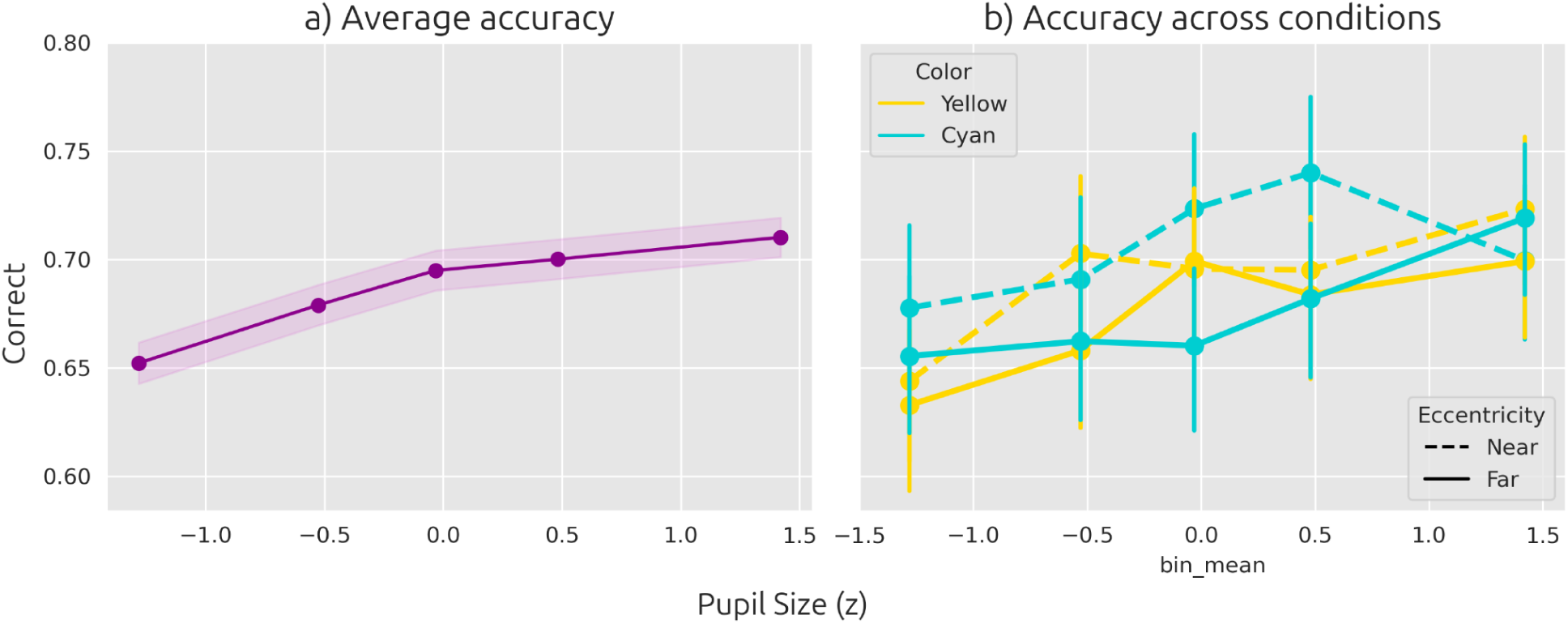
Accuracy as a function of pre-stimulus pupil size in the bright condition of Experiment 2. Pupil size is normalised within participants. Accuracy here refers to hits and correct rejections. A) Overall accuracy. The shaded area around the line represents the standard error. B) Accuracy divided into conditions. The error bars represent a bootstrap 95% confidence interval around the mean.

## Discussion

In experiment 2 we have investigated the effect of pupil size on detection of visual targets with varying colours (blue / red and yellow / cyan) and eccentricities (parafoveal and peripheral) under two different light conditions. In both light conditions we found that performance improves with increasing pupil size, replicating previous results (Eberhardt et al., 2022; Mathôt & Ivanov, 2019; Ruuskanen et al., 2025). Importantly, however, we did not observe interactions between pupil size or pupil-size squared and either colour or eccentricity, meaning that large pupils benefited detection performance regardless of these target-properties, under both light conditions.

The pattern of results in Experiment 2 is very similar to Experiment 1, and we believe that similar mechanisms are driving the results. The overall large-pupil advantage is likely to be driven by both optical effects and arousal-related effects (as defined in the Introduction). However, similarly to Experiment 1, these results do not support the proposition that pupil dilation results in a switch from rod-dominated to cone-dominated vision (Franke et al., 2022) in humans, but rather that pupil dilation globally benefits detection, regardless of which photoreceptors are (presumably) more active.

## Experiment 3

### Introduction

In Experiment 3 our main goal was to investigate the interaction between pupil size and target colour and eccentricity across a larger portion of the visual field by randomly sampling eccentricities between the near and far values used in Experiments 1 and 2. Furthermore, we were still missing a light condition that promotes mesopic vision (i.e., where both rods and cones are active) with a pure colour manipulation. Thus, in the final Experiment we tested participants in both dark and dim environments, in both cases presenting red and blue stimuli on a dark display background.

### Method

Experiment 3 consisted of two experimental sessions: a dark session and a dim session, and the same sample of participants participated in both. The order of experimental sessions was counterbalanced across participants. For both sessions, the procedure and task design was largely the same as in Experiments 1 and 2. Thus, here we will again only describe in detail those aspects that differed.

#### Participants

A total of 64 participants with normal (colour) vision participated in the experiment. However, 2 participants did not complete either session, 3 participants completed only the dark session and 1 completed only the dim session, either due to technical issues during the experiment or due to cancellations. Furthermore, some participants were excluded after pre-processing due to incomplete datafiles (likely due to unseen technical issues during recording). The final sample size used for the analysis was 54 for the dark session and 50 for the dim session. All participants were undergraduate students at the University of Groningen and received partial course credit in exchange for participation. The data was collected in the year 2024. All participants received information about the experiment and provided written informed consent prior to participation. Based on criteria developed by the EC-BSS at the University of Groningen, the study was exempt from full ethical review (study code: PSY-2324-S-0311).

#### Experimental design

##### Software, apparatus, and data acquisition

The experimental task was programmed in OpenSesame (version 4.0.13, *Melodramatic Milgram*) (Mathôt et al., 2012). Pupil data was recorded with a Gazepoint GP3 eye tracker at a sampling frequency of 60 Hz. An initial 9-point calibration procedure was performed at the beginning of the experiment and a drift correction was performed at the beginning of each trial. A chinrest was used to keep viewing distance fixed at approximately 75 cm. The experiment was conducted in either a dimly lit room (illumination as measured at the chinrest 3 LUX) or a dark room (illuminance at chinrest 0 LUX). In the dark session, participants underwent 10 minutes of dark-adaptation prior to starting the task.

##### Detection task and procedure

In both sessions, the task was presented on a black background and the targets were red and blue luminance patches (as described in Experiment 1). As opposed to the two fixed ‘near’ and ‘far’ eccentricities, this time target eccentricity could range from anywhere between parafoveal (near: 0.47 degrees of visual angle; 20 px) to peripheral (far: 9.38 degrees of visual angle; 400 px). The eccentricity was drawn randomly from this range at the beginning of every trial. The size of the targets was adjusted with a cortical magnification formula, as described in Experiment 1. The task (with only near and far eccentricities) is illustrated in Figure 5.

The task consisted of 16 practice trials and 560 experimental trials divided into 10 blocks (56 trials / block). In the first three blocks the same staircase procedure was implemented as in Experiments 1 and 2 and these trials were excluded from the analysis, leaving 392 trials.

At seven pseudo-randomly determined trials during the task (randomly drawn from predefined lists of trial numbers) participants were asked to rate their current levels of mind-wandering and focus on the task, on a 9-point Likert scale. These questions were included as part of another project, not reported here.

In addition to the detection task, participants filled in two questionnaires at the beginning of the experimental session. The first questionnaire addressed age, sex, handedness, visual correction (contact lenses, glasses or none), sleep, level of sleepiness, and consumption of nicotine, alcohol, and caffeine (in the past 10, 24, and 9 hours, respectively). The second questionnaire was the Morningness-Eveningness Questionnaire (MEQ) (Horne & Ostberg, 1976), used to measure an individual’s chronotype. After completing the questionnaires, a 5-minute resting state baseline pupil size measurement was obtained. As for the mind-wandering and focus questions during the task, the questionnaires and baseline pupil measurements were included as part of another project, the results of which will not be reported here. The questionnaire data is not included with the open data for this manuscript, but is available upon request from the corresponding author. The entire experimental session took approximately 90 minutes per participant.

##### Data processing and analysis

Initially, the same data processing and analysis pipeline was applied as in Experiments 1 and 2. However, the blink-interpolation algorithm, which was optimized for data from the EyeLink eye-tracker, introduced an artifactual trend in the data from the GazePoint eye-tracker such that the shape of the reconstructed pupil traces considerably differed from the shape of the raw traces. Thus, for both datasets of Experiment 3, blinks were not interpolated, but rather missing periods of data were simply removed. Otherwise the pre-processing pipeline was the same as for Experiments 1 and 2. For the data of both experimental sessions, the simpler model (not including a pupil squared term) had a lower AIC. Therefore, the results of this model will be reported here.

## Results

### Dark

#### Descriptive statistics

The distribution of recorded (z-scored) pupil sizes and average pupil size over time are depicted in Figure 11. On average, pupil size was larger after breaks (as indicated by dashed lines in Fig 11b) and then decreased over the course of the block. Despite this systematic variation we did not remove any trials from the analysis to avoid excluding genuine (arousal-driven) fluctuations in pupil size. However, in the Supplementary Materials we report the details of a control analysis where time on task during each block is regressed out of the pupil signal. This does not change the pattern of results, and thus, here we report the results of the analysis using the true recorded pupil size. We first examined average accuracy across the task and for each of the target colours (Table 4) and eccentricities (sorted into 5 bins; Table 5).

**Figure 11:**
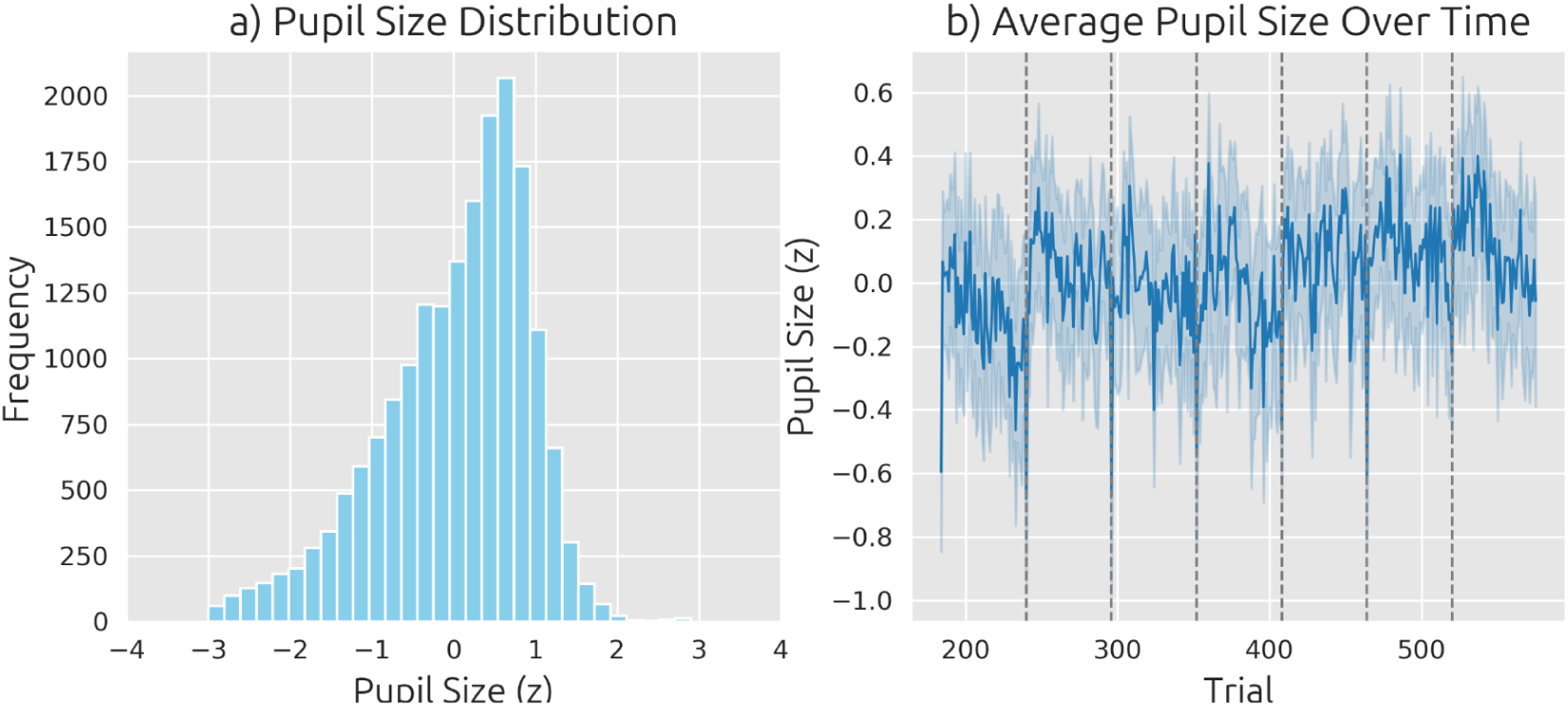
Pupil sizes recorded in the dark condition of Experiment 3 (normalised). a) Distribution of all recorded pupil sizes. b) Pupil size over time. Dashed vertical lines denote breaks. The shaded area around the line represents a bootstrap 95% confidence interval.

**Table 4.** Accuracy in the dark session of Experiment 3; divided by target colour.

| Colour | M (%) | SD |
| --- | --- | --- |
| Both | 61.33 | 0.49 |
| Blue | 59.8 | 0.49 |
| Red | 62.9 | 0.48 |

**Table 5.** Accuracy in the dark session of Experiment 3; divided by target eccentricity.

| Eccentricity<br>bin | M (%) | SD |
| --- | --- | --- |
| 0 | 49.3 | 0.50 |
| 1 | 54.01 | 0.5 |
| 2 | 62.32 | 0.48 |
| 3 | 69.18 | 0.46 |
| 4 | 71.78 | 0.45 |

#### Main analysis

Significant main effects were found for target colour (*b* = 0.27, *p* < 0.001) and eccentricity (*b* = 0.28, *p* < 0.001), whereby red targets and peripheral targets were detected better overall. There was also a significant interaction between target colour and eccentricity (*b* = -0.1, *p* < 0.001), again suggesting that red and peripheral targets are detected best. Surprisingly, the main effect of pupil size was not significant (*b* = 0.06, *p* = 0.06). However, the interaction between pupil size and eccentricity was significant (*b* = 0.03, *p* < 0.001), suggesting that larger pupils do not improve detection for parafoveally presented (i.e. near) targets. There was no significant interaction between pupil size and target colour (*b* = -0.01, *p* = 0.362). The results are depicted in Figures 12 and 13.

**Figure 12:**
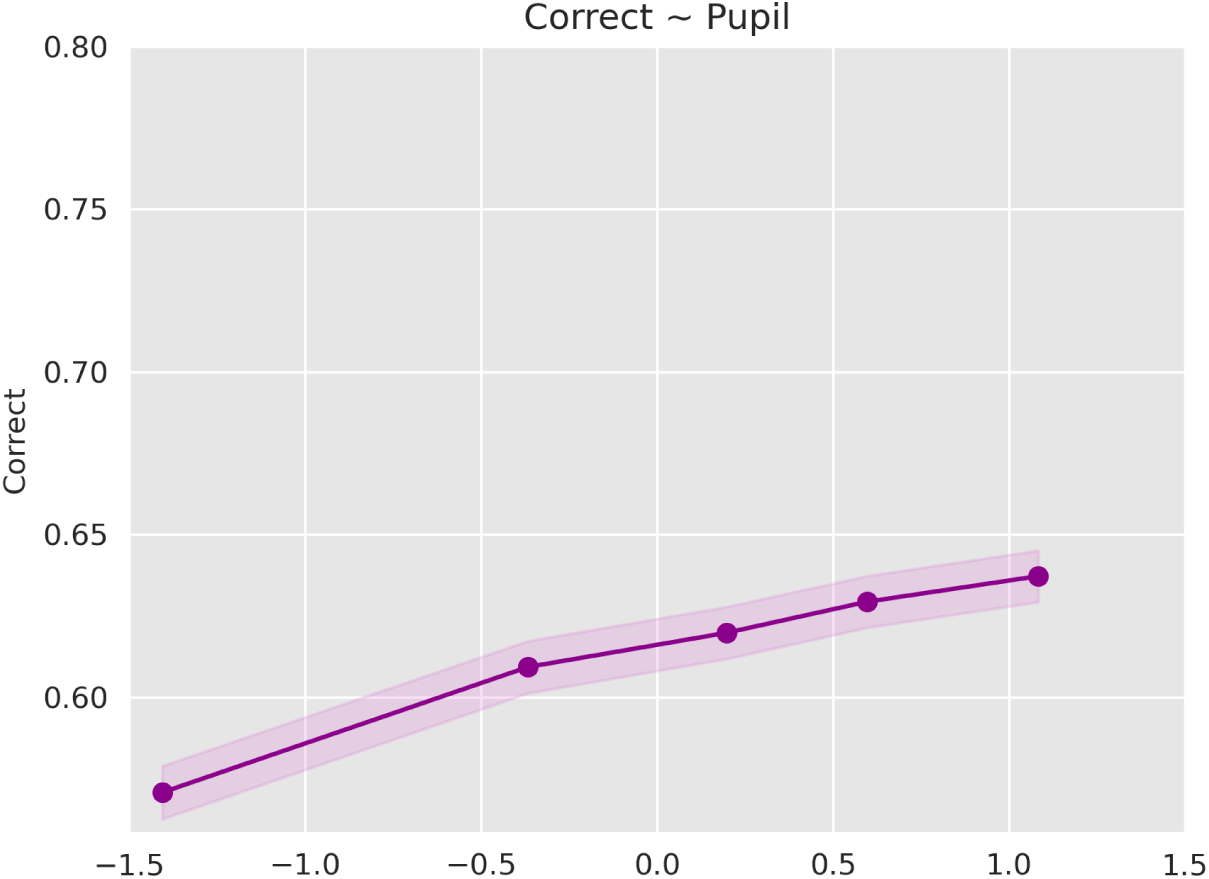
Accuracy as a function of pre-stimulus pupil size in the dark condition of Experiment 3. Pupil size is normalised within participants. Accuracy here refers to hits and correct rejections. The shaded area around the line represents the standard error.

**Figure 13:**
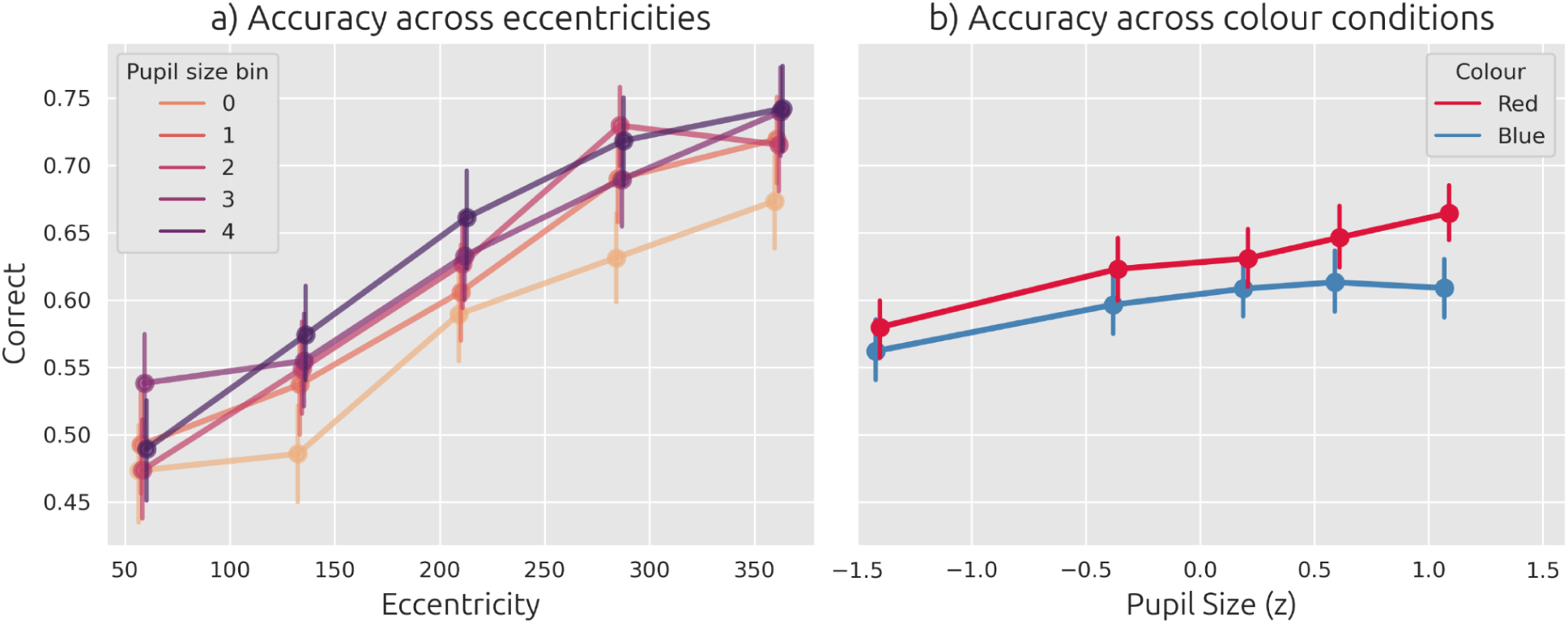
Interactions between pupil size and target eccentricity and target colour in the dark condition of Experiment 3. A) Accuracy (hits and correct rejections) as a function of eccentricity and pupil size. Lines represent pupil size bins (normalised within participants prior to binning) and eccentricity is continuous in pixels. The error bars represent a bootstrap 95% confidence interval around the mean. B) Accuracy (hits and correct rejections) as a function of colour and pupil size. The error bars represent a bootstrap 95% confidence interval around the mean.

### Dim

#### Descriptive statistics

The distribution of recorded (z-scored) pupil sizes and average pupil size over time are depicted in Figure 14. On average pupil size was larger after breaks (as indicated by dashed lines in Fig 14b) and then decreased over the course of the block. Despite this systematic variation we did not remove any trials from the analysis to avoid excluding genuine (arousal-driven) fluctuations in pupil size. However, in the Supplementary Materials we report the details of a control analysis where time on task during each block is regressed out of the pupil signal. This does not change the pattern of results, and thus, here we report the results of the analysis using the true recorded pupil size. We first examined average accuracy across the task and for each of the target colours (Table 6) and eccentricities (sorted into 5 bins; Table 7).

**Figure 14:**
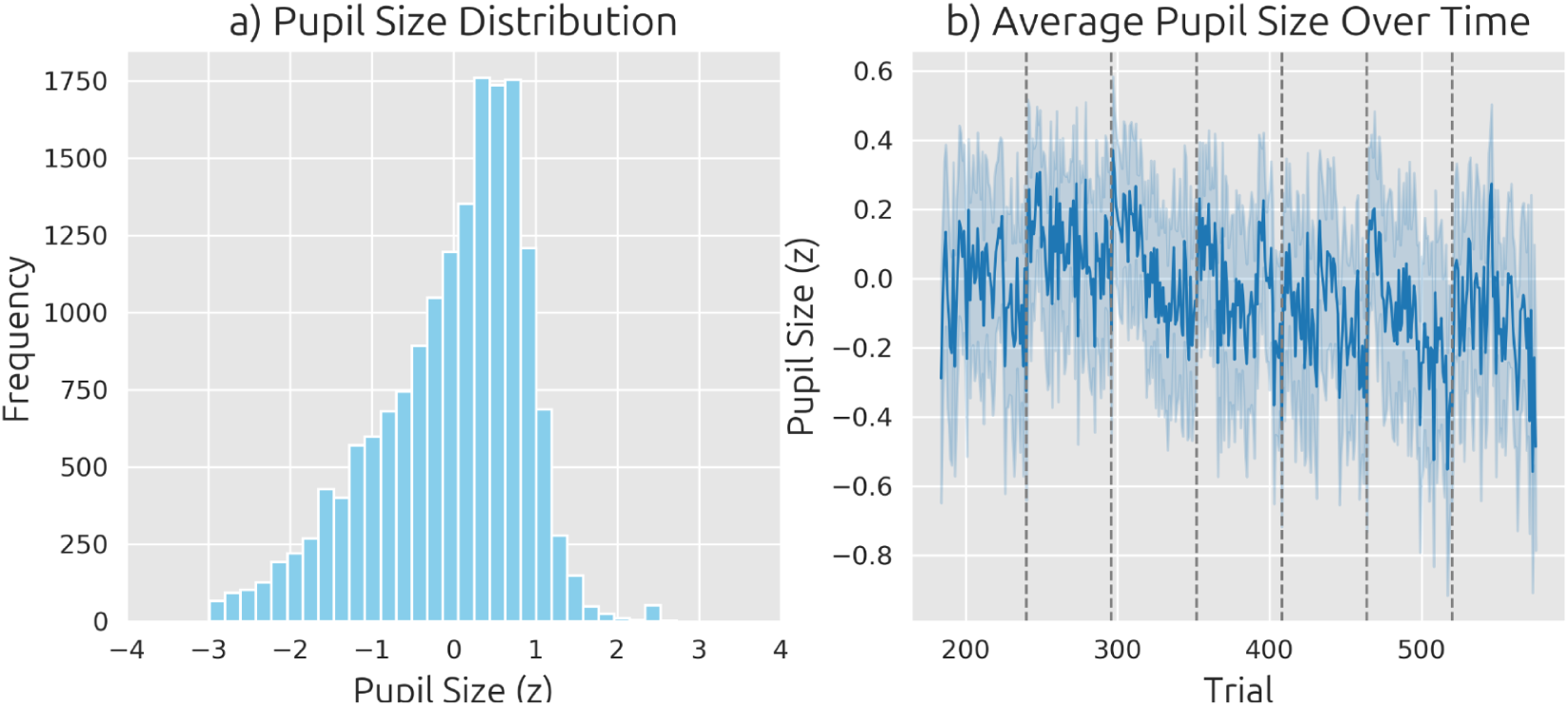
Pupil sizes recorded in the dim condition of Experiment 3 (normalised). a) Distribution of all recorded pupil sizes. b) Pupil size over time. Dashed vertical lines denote breaks. The shaded area around the line represents a bootstrap 95% confidence interval.

**Table 6.**
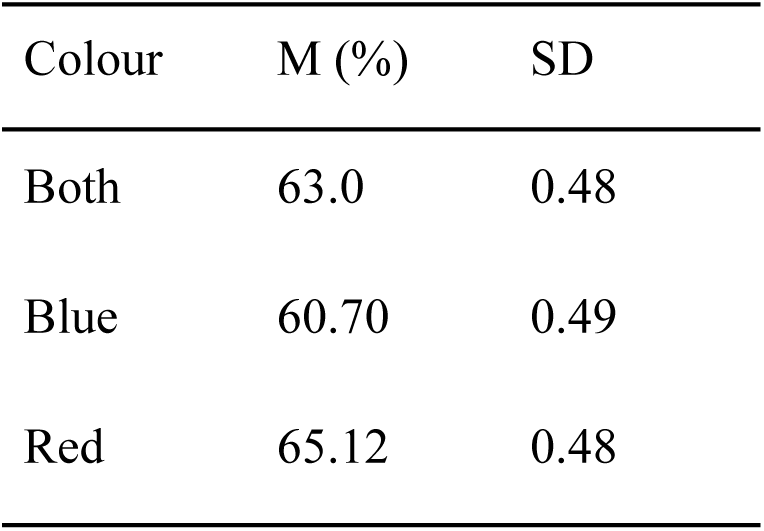
Accuracy in the dim session of Experiment 3; divided by target colour.

**Table 7.**
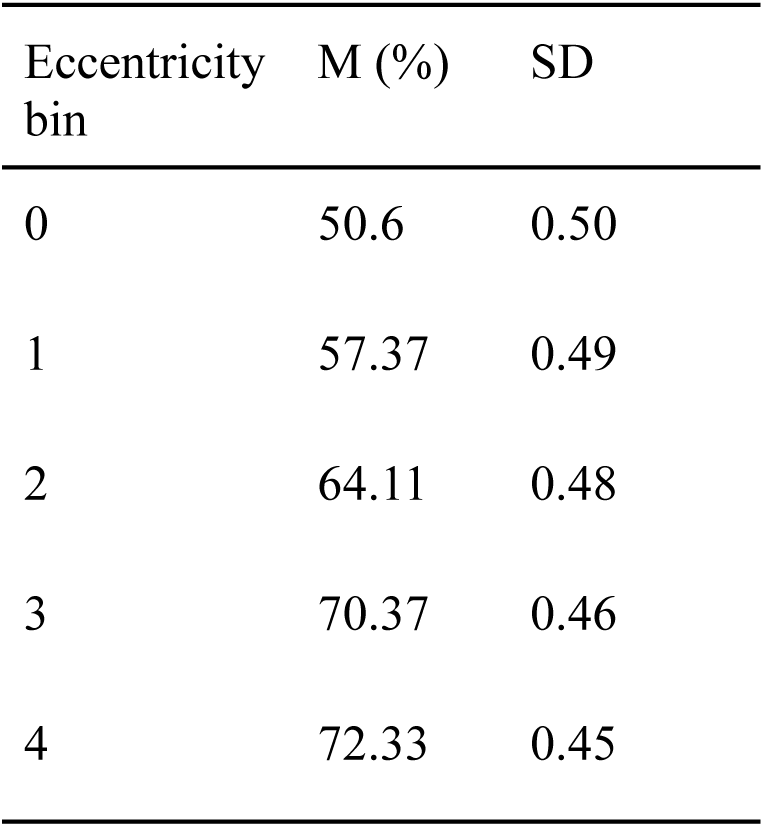
Accuracy in the dim session of Experiment 3; divided by target eccentricity.

| Eccentricity<br>bin | M (%) | SD |
| --- | --- | --- |
| 0 | 50.6 | 0.50 |
| 1 | 57.37 | 0.49 |
| 2 | 64.11 | 0.48 |
| 3 | 70.37 | 0.46 |
| 4 | 72.33 | 0.45 |

#### Main analysis

Significant main effects were found for pupil size (*b* = 0.16, *p* < .001), target colour (*b* = 0.36, *p* < .001) and eccentricity (*b* = 0.27, *p* < .001), whereby larger pupils benefited detection, and red targets and peripheral targets were detected better overall. In addition, there was a significant interaction between target colour and eccentricity (*b* = -0.13, *p* < .001). There were no significant interactions between pupil size and either colour (*b* = 0.02, *p* = .615) or eccentricity (*b* = -0.003, *p* = .851). The results are depicted in Figures 15 and 16.

**Figure 15:**
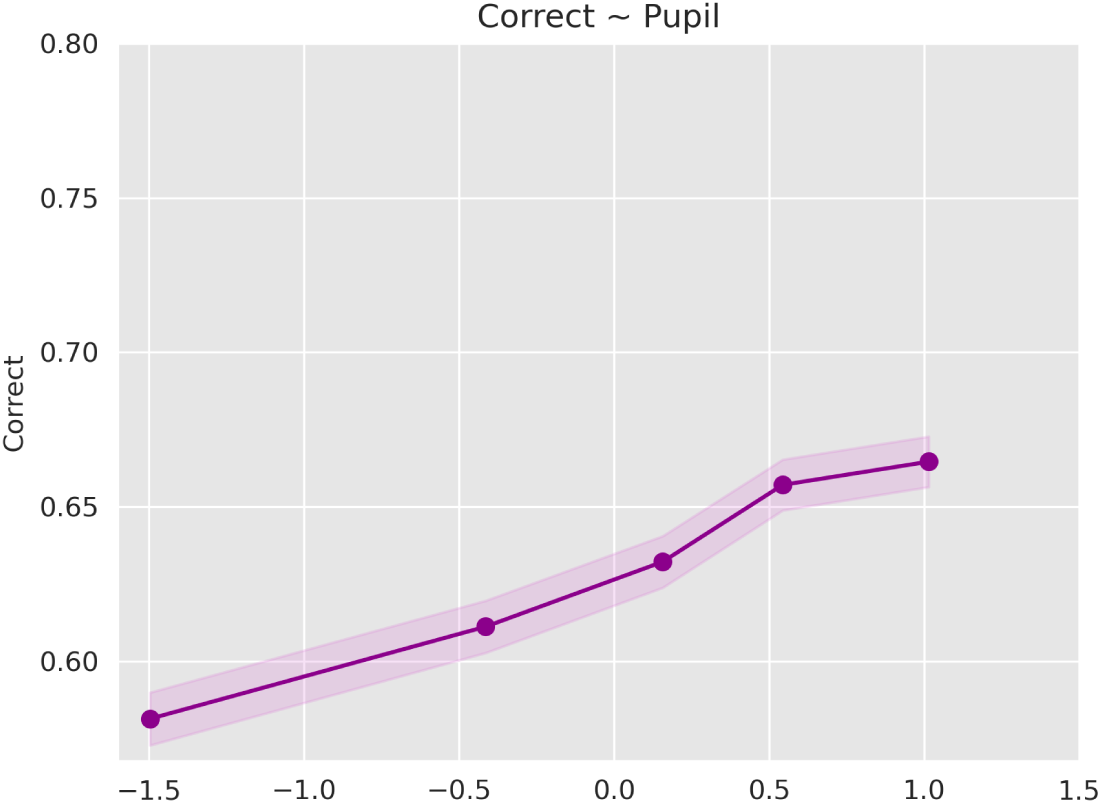
Accuracy as a function of pre-stimulus pupil size in the dim condition of Experiment 3. Pupil size is normalised within participants. Accuracy here refers to hits and correct rejections. The shaded area around the line represents the standard error.

**Figure 16:**
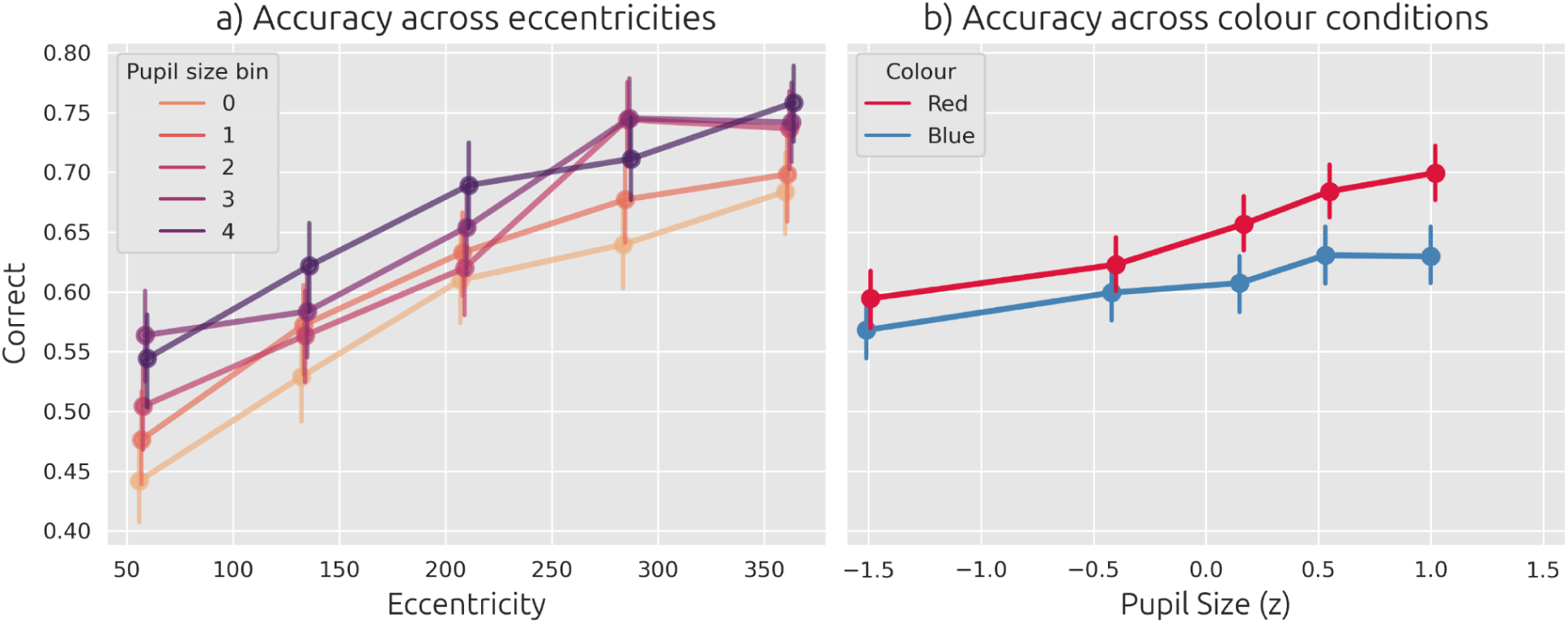
Interactions between pupil size and target eccentricity and target colour in the dim condition of Experiment 3. A) Accuracy (hits and correct rejections) as a function of eccentricity and pupil size. Lines represent pupil size bins (normalised within participants prior to binning) and eccentricity is continuous in pixels. The error bars represent a bootstrap 95% confidence interval around the mean. B) Accuracy (hits and correct rejections) as a function of colour and pupil size. The error bars represent a bootstrap 95% confidence interval around the mean.

## Discussion

In experiment 3 we have investigated the effect of pupil size on detection of near-threshold visual targets with varying colours (blue and red), at randomly sampled (as opposed to fixed) eccentricities, under two different light conditions. In the dim condition we found that performance improves with increasing pupil size, replicating previous results (Eberhardt et al., 2022; Mathôt & Ivanov, 2019; Ruuskanen et al., 2025). In the dark condition this effect was not significant. However, in the dark condition we did observe an interaction between pupil size and target eccentricity, suggesting that larger pupils improve detection more at more peripheral eccentricities compared to the nearest eccentricity. In the dim condition no interactions between pupil size and target properties were observed. Furthermore (although not of theoretical importance), in both light-conditions detection accuracy increased when targets were presented more peripherally, and red targets were detected better than blue ones. These results will be further discussed below, in the General Discussion.

## General discussion

Here we have presented the results of three experiments designed to investigate how pupil size interacts with target colour, target eccentricity, and retinal adaptation-state in determining near-threshold detection performance. In all but one out of the five collected datasets we have shown that larger pupils benefit detection, regardless of the properties of the targets or the adaptation-state of the retina. Furthermore, in the dataset where the general relationship was not present, we did show a significant interaction between pupil size and target eccentricity, suggesting that in this dataset the effect of pupil size on detection differed at different eccentricities. These findings replicate and extend those of previous research (Eberhardt et al., 2022; Mathôt & Ivanov, 2019; Ruuskanen et al., 2025) by showing that the robust relationship between pupil size and near-threshold visual detection performance holds for different stimulus colours, eccentricities, and retinal-adaptation states.

One underlying motivation for the experiments presented here was investigating the possibility that a change in pupil size could induce a shift from rod-dominated to cone-dominated vision, following findings from research in mice (Franke et al., 2022). To this end, we used combinations of stimuli that differentially activate either the rods or the cones, as well as conducted experiments under varying light conditions to promote reliance on either scotopic, mesopic, or photopic vision. However, we did not observe any reliable interactions between target-properties and pupil size, or different patterns of results under different light conditions. Thus, we conclude that in humans, the increased light influx resulting from pupil dilation is not enough to induce a switch from rod to cone-dominance. It is, however, sufficient to improve visual detection sensitivity, presumably due to an increased signal-to-noise ratio.

As discussed in the Introduction, we propose that the general relationship between pupil size and near-threshold detection is a reflection of an optical effect and an arousal effect. The optical effect refers to the improvement in visual sensitivity that results from a dilated pupil allowing more light on the retina (Ruuskanen et al., 2025; Vilotijević & Mathôt, 2023). The arousal effect refers to the increased visual sensitivity that results from a more optimal level of arousal, as suggested by the classical Yerkes-Dodson curve of performance as a function of arousal, whereby performance is best at medium levels of arousal, and deteriorates at low and high arousal-states (Yerkes & Dodson, 1908). Therefore, these effects also differ in their visual representation (i.e., when plotting accuracy as a function of pupil size). The optical effect should visually roughly resemble a linear relationship, with accuracy steadily increasing with increasing pupil size. The arousal effect on the other hand should visually resemble an inverted-U-shape, following the Yerkes-Dodson curve. A combination of both thus results in a graph with an S-shaped or slightly curved line, where larger pupils are generally associated with improved detection, but the relationship is not strictly linear.

The results presented here roughly align with this proposition, whereby there is some evidence for both mechanisms. A visual inspection of the general relationship between pupil size and accuracy in all datasets (Figure 1) suggests that both linear and non-linear effects are present; that is, the most notable improvement in accuracy is evident between small and medium pupil sizes, with little to no improvement from medium to large pupil sizes. However, when performing a statistical model comparison procedure, a simple model with only a linear component was favoured over a complex model with an additional quadratic component in all analyses. Thus, we suggest that in primarily visual tasks where required cognitive effort is low (such as the task used here), the optical effect is more important in determining performance than the arousal effect. This interpretation is supported by evidence showing that in the same task the effects of pupil size and EEG measures related to arousal exert partly independent influences on performance (Ruuskanen et al., 2025).

However, this general pattern of results differed slightly in the dark condition in Experiment 3, where participants were dark adapted and assumed to operate under scotopic vision. In this condition the general relationship between pupil size and accuracy was not reliably present (although there was a trend) but there was a significant interaction between pupil size and eccentricity; at the nearest eccentricity there were no differences in performance between different pupil size bins, while at further eccentricities larger pupils were associated with higher accuracy. Furthermore, accuracy was generally low (45-50%) at the nearest eccentricity. These findings align with previous research on visual search under scotopic light conditions, where only rods are active (Paulun et al., 2015). Specifically, when the cones are not active, vision at or near the fovea deteriorates (Barbur & Stockman, 2010; Paulun et al., 2015). Here, this resulted in low performance at the nearest eccentricity, with no large-pupil advantage.

Notably, our general pattern of results differs quite substantially from several other studies investigating the relationship between pupil size and performance on near-threshold perceptual tasks. While we have consistently observed a pattern that is more accurately described as linear rather than inverted-U shaped (see also; Ruuskanen et al., 2025), several other studies show the opposite pattern (e.g., Beerendonk et al., 2024; de Gee et al., 2024; McGinley et al., 2015; Podvalny et al., 2021; Schriver et al., 2018). We believe there are three main reasons for this discrepancy. Firstly, the type of the stimulus matters. As mentioned above, we believe that the optical effect of pupil size on performance is especially relevant in visual tasks; in other types of tasks, such as auditory detection (de Gee et al., 2024; McGinley et al., 2015) and tactile discrimination (Schriver et al., 2018), which do not benefit from enhanced *visual* sensitivity, task performance is likely to be mainly driven by fluctuations in arousal, manifesting as an inverted-U shaped relationship between pupil size and performance. Secondly, the outcome measure seems to matter; in one study on visual near-threshold detection a linear relationship between pupil size and hit rate was observed, while the relationship between pupil size and sensitivity (d’) was inverted-U shaped (Podvalny et al., 2021). Similarly, other studies investigating accuracy or hit rate, as opposed to sensitivity, seem to find more linear patterns (Eberhardt et al., 2022; Mathôt & Ivanov, 2019). Finally, the range of the arousal fluctuations measured could influence results; as noted by Beerendonk and colleagues (2024) it is likely that the fluctuations observed here only cover low-to-medium levels of arousal (since the task is not especially engaging), thus leading to observing a linear relationship, and potentially, an emphasis on optical effects over arousal effects.

In conclusion, here we have shown that larger pre-stimulus pupil size is associated with higher accuracy in near-threshold visual detection and that this general relationship is not modulated by target colour, eccentricity, or retinal adaptation-state. These results align with findings from previous research (Eberhardt et al., 2022; Mathôt & Ivanov, 2019; Ruuskanen et al., 2025) and further expand on them by showing that the large-pupil advantage for detection is not only present for black and white peripherally presented targets. More generally, our findings add to the growing body of research on the unique effects of pupil size on visual perception and visually guided behaviour.

## Acknowledgments

We would like to thank the following students for their valuable help with data collection: Ward Eiling, Daria Weiden, Elizabeta Chiriac, Tessa Mesken, Maud Hooijman, Hannah Postma, and Louise Völker.

## Funding

This research was funded by the Dutch Research Council (NWO; Grant VI.Vidi.191.045).

## Notes

### Competing Interest Statement

The authors have declared no competing interest.

### Summary of Updates

Added supplementary materials with additional analyses

