## Supplementary materials for "Pupil size correlates with near-threshold detection performance irrespective of stimulus colour, eccentricity, or retinal adaptation-state"

Supplementary Methods and Results associated with Ruuskanen & Mathôt: Pupil size correlates with near-threshold detection performance irrespective of stimulus colour, eccentricity, or retinal adaptation-state

### Control analysis 1: inspecting the effect of target location

To control for target location beyond eccentricity, we conducted an additional analysis with target location (specified as the quadrant the target was presented in: upper left, lower left, lower right and lower left) as a factor. First, we verified that targets were presented approximately equally in all four quadrants (i.e., ~25% of targets presented in each). This was the case for all experiments. Next, on each dataset, we fitted a Generalised Mixed Linear Model (GLM) predicting accuracy from pre-stimulus pupil size and target location, as well as their interaction. We also fitted subject-specific intercepts and slopes for the main effects. Target colour and eccentricity were not included in the model. The results of the GLM are reported below.

#### Experiment 1

The GLM showed a significant main effect of pupil size ($b = 0.28$, $p < .001$). Furthermore, pupil size significantly interacted with target location, such that larger pupils did not improve accuracy on the left side of the lower visual field ($b = -0.15$, $p < .001$) to the same extent as in other locations. The relationship between pupil size and accuracy at different target locations in Experiment 1 is depicted in Figure 1.

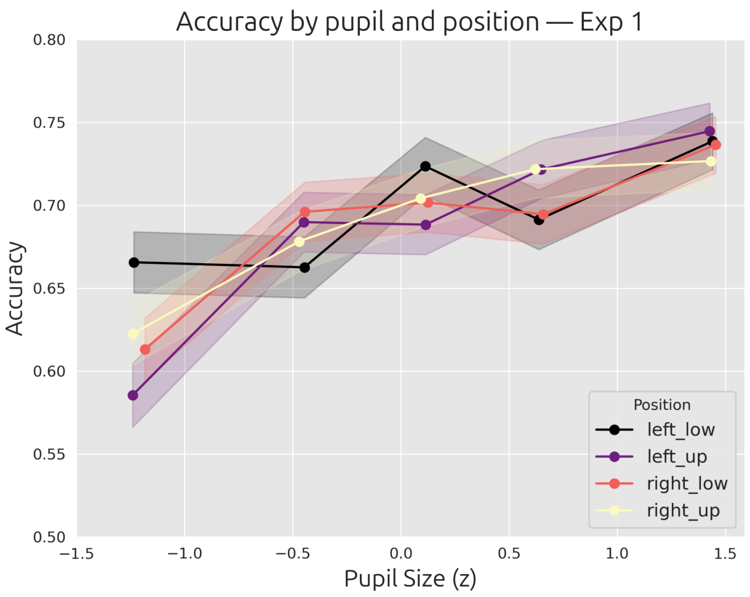


*Figure 1:* Accuracy at different target locations as a function of z-scored pupil size (Experiment 1). The shaded area around the line represents the standard error.

### Experiment 2

#### *Bright condition*

The GLM showed a significant main effect of pupil size ($b = 0.13$, $p = .002$) such that larger pupils improved detection, and of target location, such that targets in the lower visual field were associated with decreased accuracy both on the left ($b = -0.41$, $p < .001$) and on the right ($b = -0.43$, $p < .001$). There were no interactions between pupil size and target location. The relationship between pupil size and accuracy at different target locations is depicted in Figure 2a.

#### *Dark condition*

The GLM showed significant main effects of pupil size, such that larger pupils were associated with improved accuracy ($b = 0.13$, $p = .005$), and of target location, such that targets in the lower visual field (right: $b = 0.82$, $p < .001$; left: $b = 0.75$, $p < .001$) and in the upper right visual field ($b = 0.14$, $p = .007$) were associated with better accuracy than those in the upper left visual field. There were no interactions between pupil size and target location. The relationship between pupil size and accuracy at different target locations is depicted in Figure 2b.

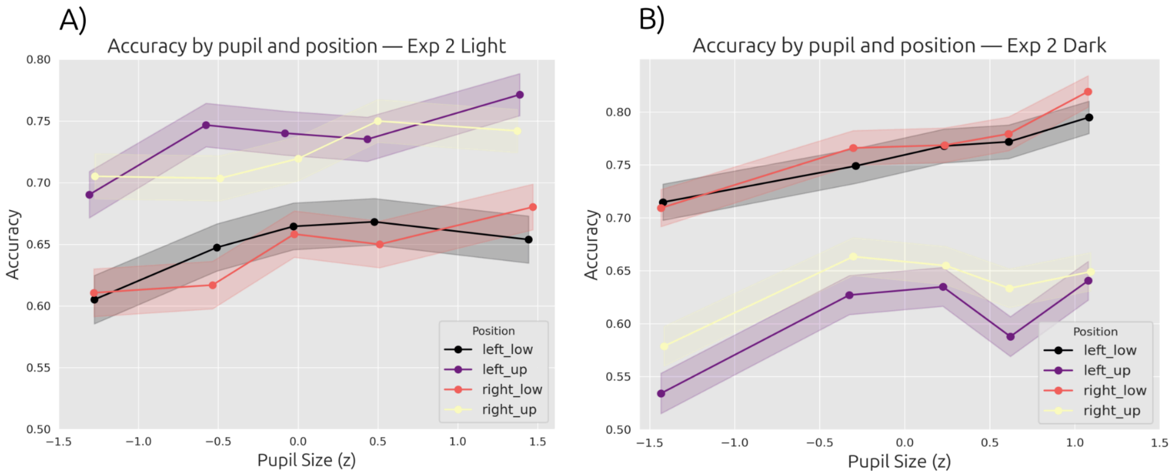


*Figure 2:* Accuracy at different target locations as a function of z-scored pupil size (Experiment 2). The shaded area around the line represents the standard error. A) The light condition of Experiment 2. B) The dark condition of experiment 2.

### Experiment 3

#### *Dim condition*

The GLM showed significant main effects of pupil size, such that larger pupils were associated with improved accuracy ($b = 0.11$, $p = .008$), and of target location, such that targets in the lower visual field were associated with improved accuracy both on the left ($b = 0.76$, $p < .001$) and on the right ($b = 0.85$, $p < .001$). There were no interactions between pupil size and target location.  The relationship between pupil size and accuracy at different target locations is depicted in Figure 3a.

#### *Dark condition*

The GLM showed significant main effects of pupil size, such that larger pupils were associated with improved accuracy ($b = 0.09$, $p = .011$), and of target location, such that targets in the lower visual field were associated with improved accuracy both on the left ($b = 0.74$, $p < .001$) and on the right ($b = 0.75$, $p < .001$). There were no interactions between pupil size and target location. The relationship between pupil size and accuracy at different target locations is depicted in Figure 3b.

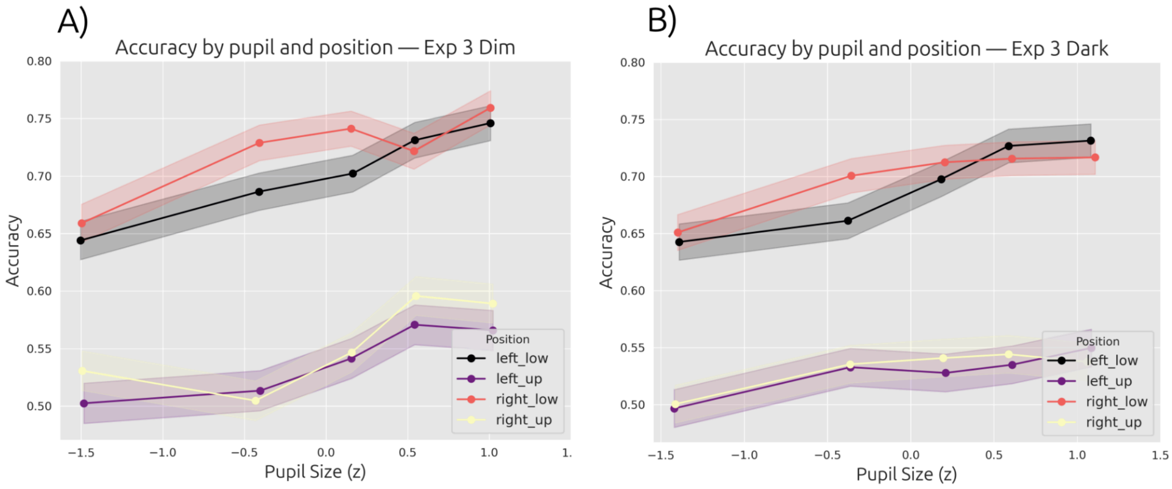


*Figure 3:* Accuracy at different target locations as a function of z-scored pupil size (Experiment 3). The shaded area around the line represents the standard error. A) The dim condition of Experiment 3. B) The dark condition of experiment 3.

### Control analysis 2: time on task

To control for potential time-on-task effects we repeated our main analysis after removing the effect of time from pre-stimulus pupil size. Specifically, we first fit linear and quadratic regression models predicting pupil size from trial number within a block, for each experiment separately. We then compared the AIC of each model (or BIC if AIC was identical) to determine which one fit the data better. For all experiments the quadratic model had better fit. We then fitted the chosen model for each participant separately, and recorded the residuals. Next, we repeated our GLM analyses as described in the Manuscript, but replaced the original pupil variable with the residuals. This did not change the results. Details are reported below.

### Experiment 1

The GLM showed significant main effects of residual pupil size ($b = 0.22$, $p < .001$), whereby larger pupils improved detection, and of eccentricity ($b = 0.13$, $p = .009$), whereby peripheral targets were detected better. Further, there was a significant interaction between colour and eccentricity ($b = 0.08$, $p < .001$), whereby red and peripheral targets were detected best. This pattern is consistent with the original analysis.

### Experiment 2

#### *Dark condition*

The GLM showed a significant main effect of pupil size, such that larger pupils were associated with improved accuracy ($b = .14$, $p < .001$). This is consistent with the original analysis. Furthermore, there was a significant interaction between target colour and eccentricity ($b = .04$, $p = .036$)

#### *Bright condition*

The GLM showed a significant main effect of pupil size ($b = 0.06$, $p = .011$), whereby larger pupils improved detection. This is consistent with the original analysis. There were no other significant effects.

### Experiment 3

#### *Dark condition*

The GLM showed significant main effects for target colour ($b = 0.36$, $p < .001$) and eccentricity ($b = 0.28$, $p < .001$). There was also a significant interaction between target colour and eccentricity ($b = -0.1$, $p < .001$). The main effect of pupil size was not significant ($b = 0.04$, $p = .429$). However, the interaction between pupil size and eccentricity was significant ($b = 0.03$, $p$

= .039), suggesting that larger pupils do not improve detection for parafoveally presented (i.e. near) targets. This pattern is consistent with the original analysis.

***Dim condition***

The GLM showed significant main effects of pupil size ($b = 0.16$, $p = .001$), target colour ($b = 0.49$, $p < .001$) and target eccentricity (b = 0.27, p < .001). There was also a significant interaction between colour and eccentricity (b = -0.14, p < .001). This pattern is consistent with the original analysis.

### Pupil size and other variables

#### Signal detection theory

According to signal detection theory (SDT), accuracy in detection type tasks is a combination of the observer’s sensitivity (d’) and a decision criterion (i.e., the level of evidence needed to report that a signal is present). Figures 4 and 5 depict the relationship between pre-stimulus pupil size and d’ and criterion, respectively. In this visualisation, pupil size is binned into 5 bins across participants, and the aggregate measures are computed for each bin. The figures indicate that larger pupils are generally associated with improved sensitivity and a more liberal criterion, in all experiments. The shape of the relationship between pupil size and d’ aligns with our results on accuracy, suggesting that the effect of pupil size on accuracy may reflect increased sensitivity at larger pupil sizes.

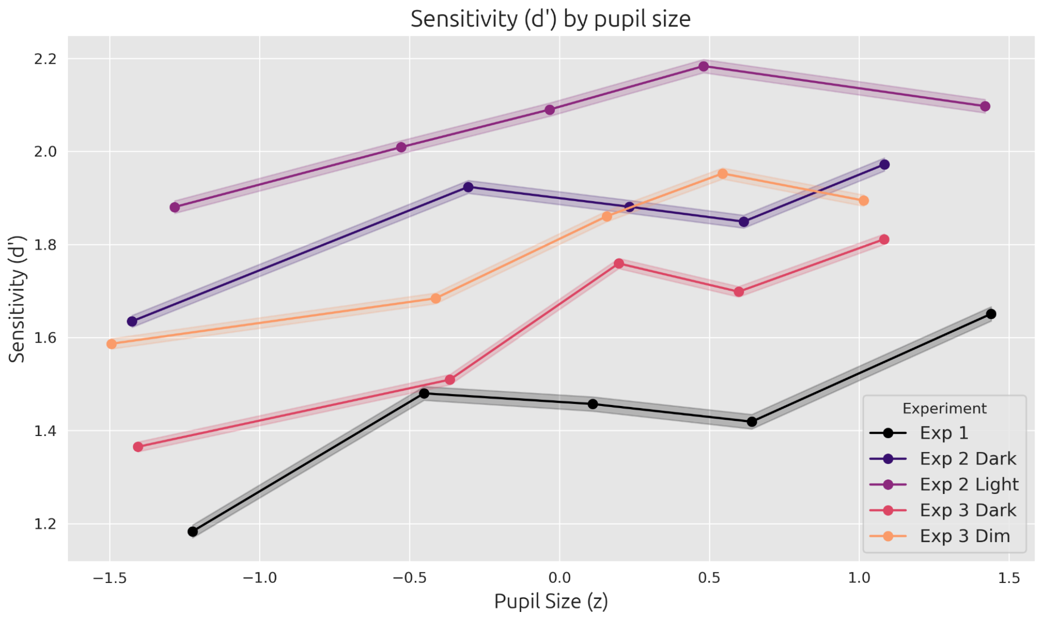


*Figure 4:* Sensitivity as a function of z-scored pupil size, across experiments. The shaded area around the lines represents the standard error.

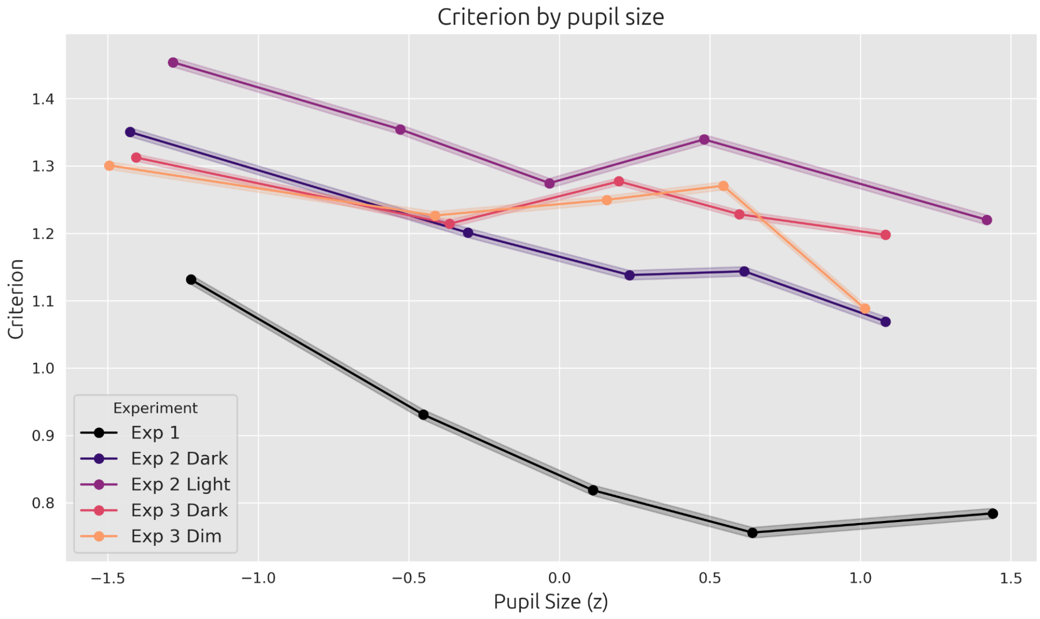


*Figure 5:* Criterion as a function of z-scored pupil size, across experiments. The shaded area around the lines represents the standard error.

**Reaction time on hit trials**

To investigate the potential relationship between pupil size and reaction time (RT), we fitted a linear mixed effects model predicting RT from pre-stimulus pupil size, with random intercepts for subjects. The model was fit separately for each experiment. Prior to the analysis we removed implausible (i.e., too short: less than 200 ms, or too long: more than 1400 ms) reaction times. The results for each experiment are described below.

***Experiment 1***

The GLM showed a significant main effect of pupil size, such that larger pupils were associated with faster reaction times ($b$ = -14.08, $p$ < .001; Figure 6).

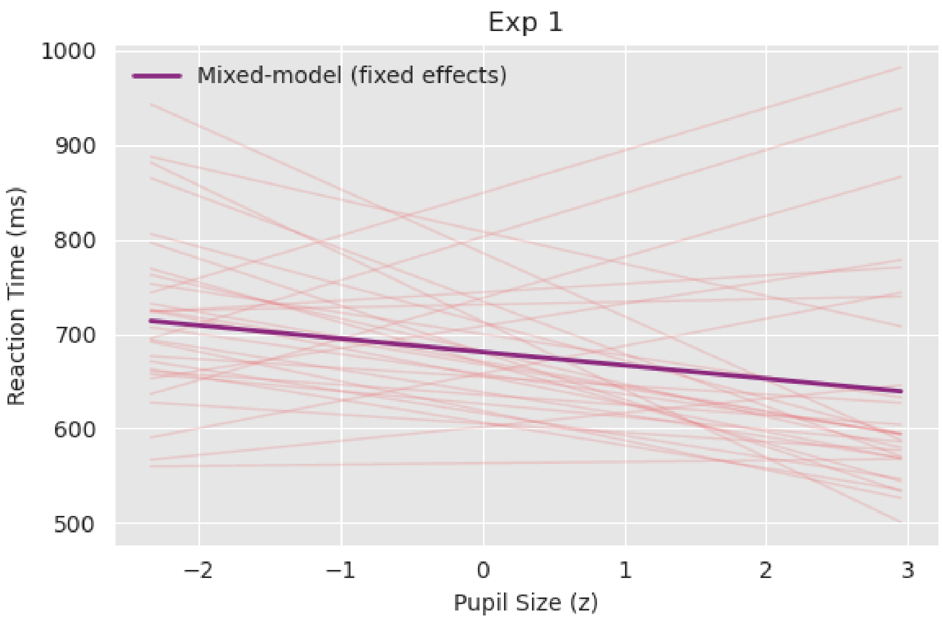


*Figure 6:* The relationship between pupil size and reaction time (RT) on hit trials in Experiment 1. Lightly coloured lines represent individual participants and the darker line the overall slope of the GLM.

***Experiment 2***

**Dark condition**

The effect of pupil size on RT was not significant ($b$ = -0.35, $p$ = .728; Figure 7a)

**Bright condition**

The GLM showed a significant main effect of pupil size, such that larger pupils were associated with faster reaction times (*b* = -11.38, *p* = .003; Figure 7b).

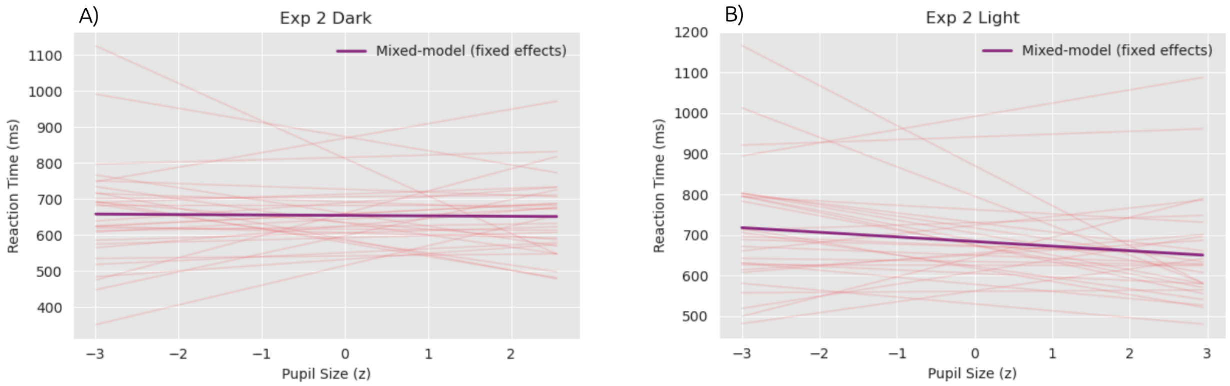


*Figure 7:* The relationship between pupil size and reaction time (RT) on hit trials in Experiment 2. Lightly coloured lines represent individual participants and the darker line the overall slope of the GLM. A) Dark condition. B) Bright condition.

***Experiment 3***

**Dark condition**

The GLM showed a significant main effect of pupil size, such that larger pupils were associated with faster reaction times (*b* = -9.2, *p* = .003; Figure 8a).

**Dim condition**

The GLM showed a significant main effect of pupil size, such that larger pupils were associated with faster reaction times (*b* = -7.16, *p* = .016; Figure 8b).

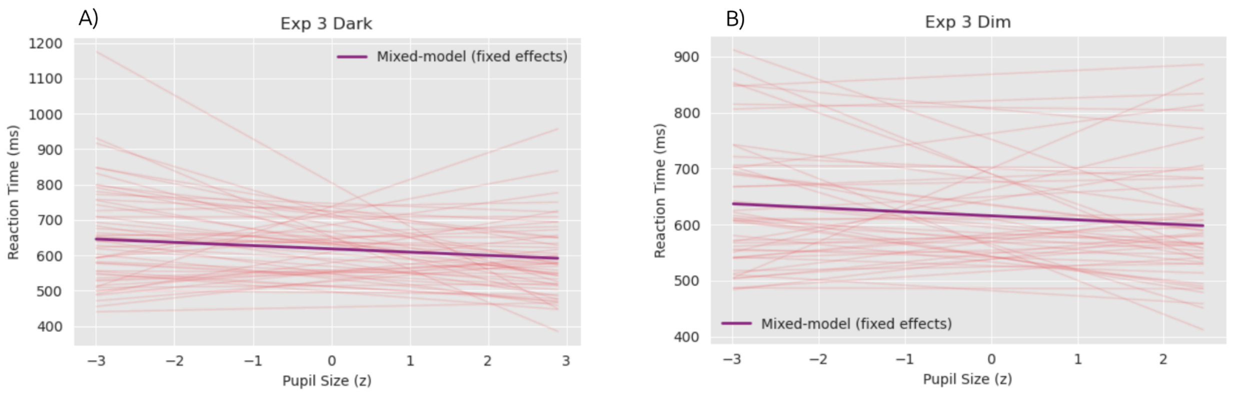


*Figure 8:* The relationship between pupil size and reaction time (RT) on hit trials in Experiment 3. Lightly coloured lines represent individual participants and the darker line the overall slope of the GLM. A) Dark condition. B) Dim condition.

### Blinks and saccades

To characterize the relationship between pupil size and blink rate, we counted the number of blinks on each trial (using the lists of blink start and times as recorded by the eyetracker). Then we binned pupil size into five equal-sized bins and computed the average number of blinks in each bin. These are depicted in Figure 9. This analysis was only conducted for experiments 1 and 2 since the eyetracking data of Experiment 3 was collected with a Gazepoint eyetracker which does not provide blink start and end times.

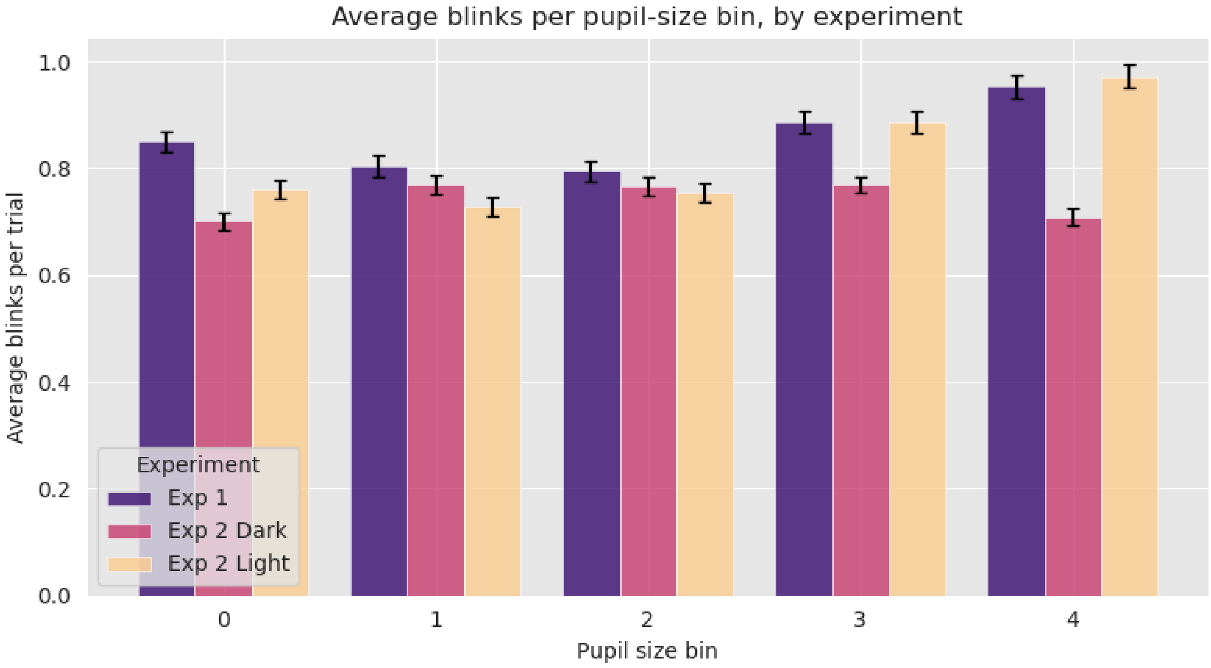


*Figure 9:* Average number of blinks per trial across pupil size bins in experiments 1 and 2. The caps on the bars represent the standard error of the mean. Differently coloured bars represent different experiments.

Since participants were required to maintain fixation throughout the task, saccades were determined as deviations from the centre that exceeded 3° of visual angle and lasted for 10 or more consecutive samples. These were computed using the lists of x and y coordinates recorded by the eyetracker. Most trials did not contain any saccades: for a given participant on average <5% of the trials in Experiment 1, and <10% and <5% of the trials in the two conditions of Experiment 2 (dark and bright, respectively) contained one. Therefore, instead of counting the number of saccades as we did for blinks, we simply recorded whether a saccade was present on a trial or not, and whether it happened prior to or after the presentation of a target. Trials with pre-target saccades were removed. Figure 10 shows the probability of a saccade *after* the target as a function of pupil size. This analysis was only conducted for experiments 1 and 2 since the eyetracking data of Experiment 3 was collected with a Gazepoint eyetracker which records eye position in arbitrary units that we could not convert to pixel coordinates.

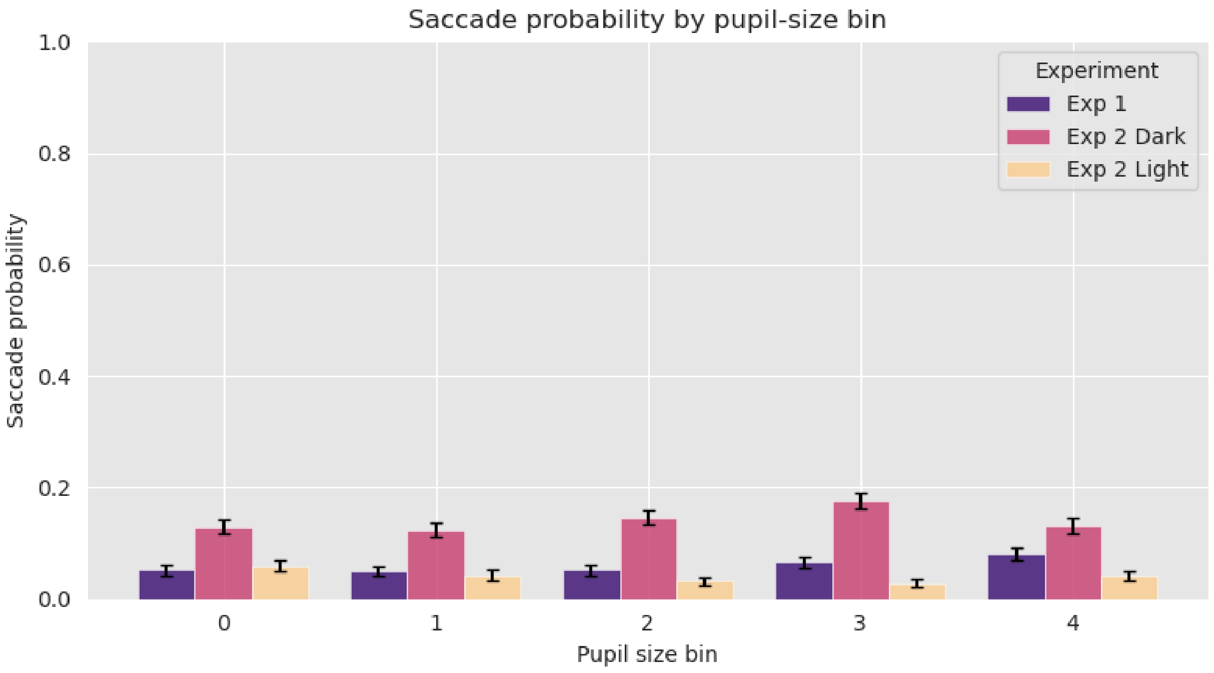


*Figure 10:* Saccade probability after target presentation across pupil size bins in experiments 1 and 2. The caps on the bars represent the standard error of the mean. Differently coloured bars represent different experiments.

### Additional Figures

Figures 11-16 below depict the average pupil size time course locked to stimulus presentation (or the moment a stimulus would have been presented on target-absent trials), separately for all trials and hit trials (i.e., trials where a stimulus was present and a response was given). Generally, as expected, on hit trials the pupil dilates slightly due to the combination of target detection and the associated button press. However, given that hits are relatively uncommon due to the difficulty of the task, this trend is not present in the average trace of all trials. The highlighted area of the figure was used in all analyses reported both in the Manuscript and here.

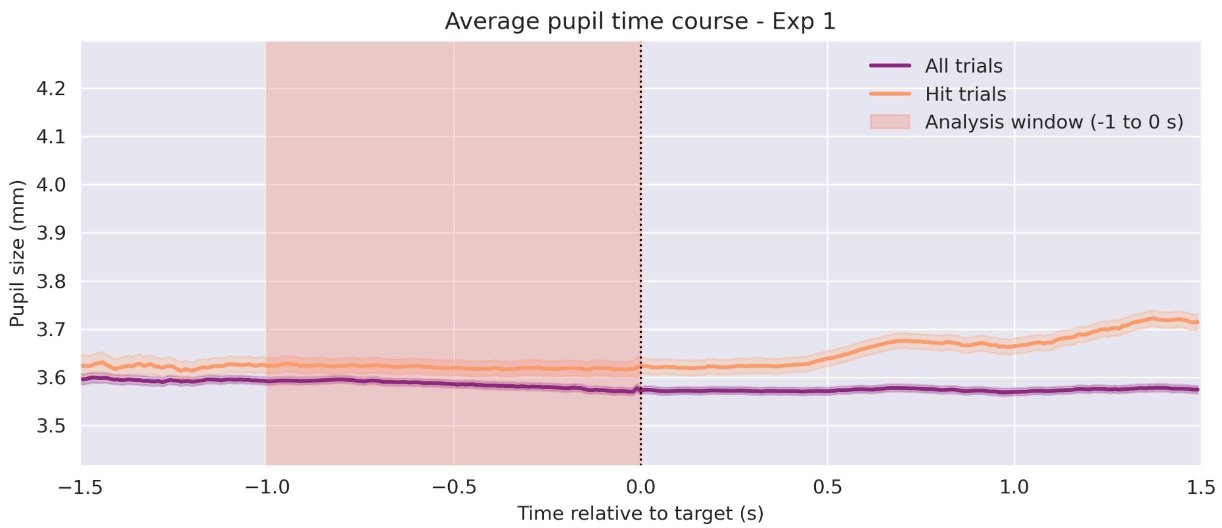


*Figure 11:* Average pupil time course throughout all trials (purple) and hit trials (orange) in Experiment 1. The highlighted area corresponds to the analysis window. *Note*: the slight discontinuity at time 0 results from concatenating two traces that preprocessed independently.

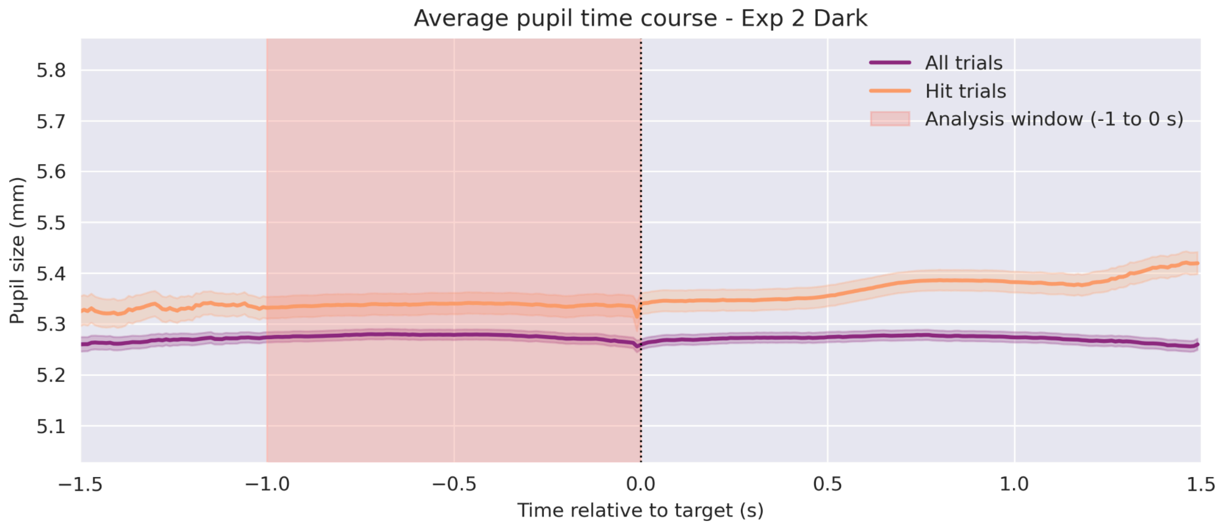

*Figure 12:* Average pupil time course throughout all trials (purple) and hit trials (orange) in Experiment 2, dark condition. The highlighted area corresponds to the analysis window. *Note*: the slight discontinuity at time 0 results from concatenating two traces that preprocessed independently.

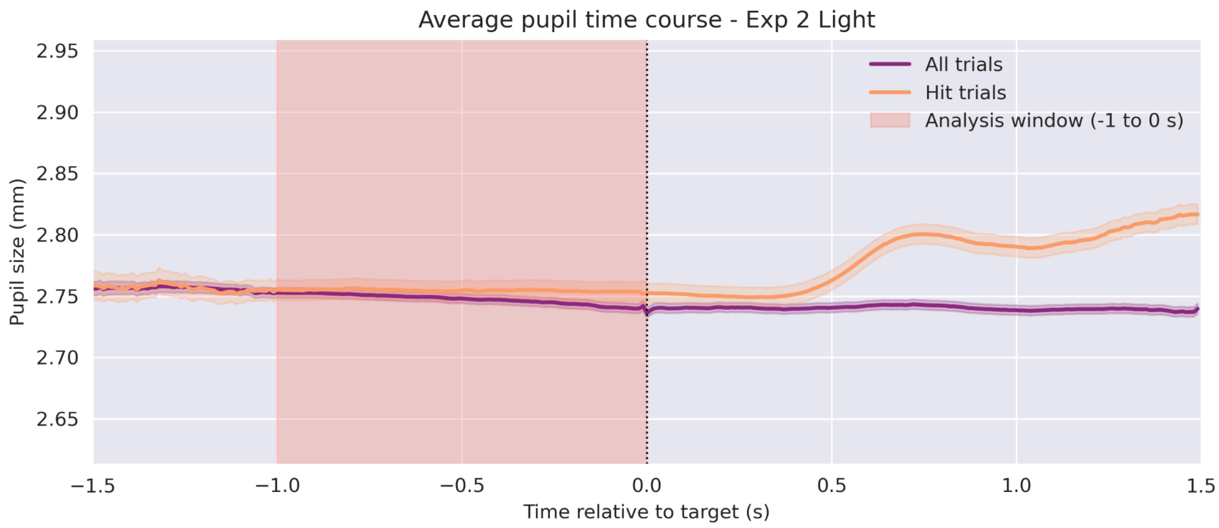


*Figure 13:* Average pupil time course throughout all trials (purple) and hit trials (orange) in Experiment 2, light condition. The highlighted area corresponds to the analysis window. Note: the slight discontinuity at time 0 results from concatenating two traces that preprocessed independently.

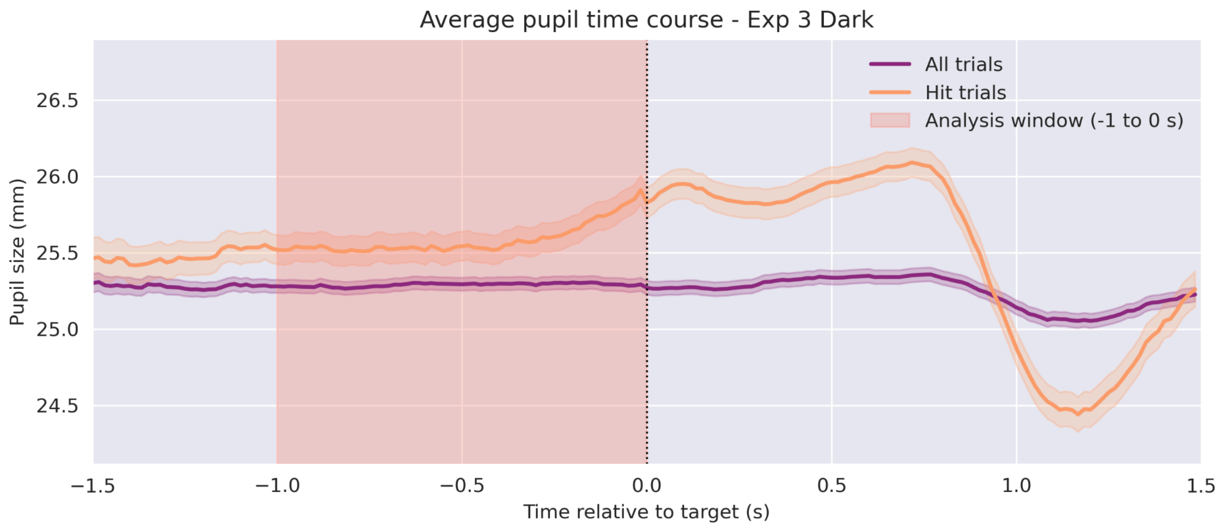


*Figure 14:* Average pupil time course throughout all trials (purple) and hit trials (orange) in Experiment 3, dark condition. The highlighted area corresponds to the analysis window. *Note 1*: pupil

size in arbitrary units as recorded by the Gazepoint eyetracker. *Note 2*: the slight discontinuity at time 0 results from concatenating two traces that preprocessed independently.

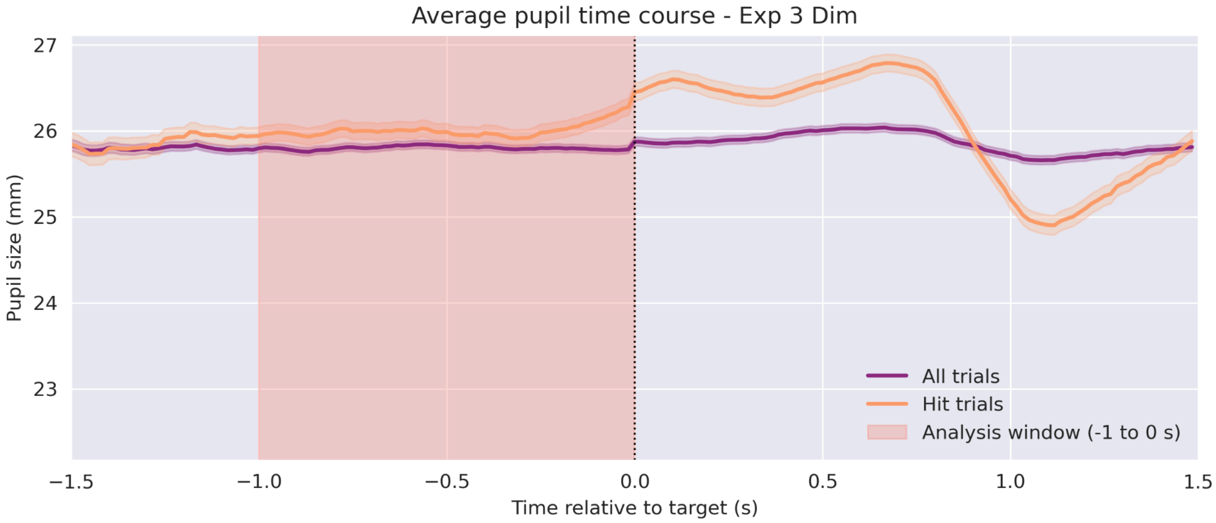


*Figure 15:* Average pupil time course throughout all trials (purple) and hit trials (orange) in Experiment 3, dim condition. The highlighted area corresponds to the analysis window. *Note 1*: pupil size in arbitrary units as recorded by the Gazepoint eyetracker. *Note 2*: the slight discontinuity at time 0 results from concatenating two traces that preprocessed independently.
